# GCNA interacts with Spartan and Topoisomerase II to regulate genome stability

**DOI:** 10.1101/570200

**Authors:** Gregory M. Davis, Gregoriy A. Dokshin, Ashley D. Sawle, Matthew D. Eldridge, Katherine A. Romer, Taylin E. Gourley, Luke W. Molesworth, Hannah R. Tatnell, Ahmet R. Ozturk, Dirk G. de Rooij, Gregory J. Hannon, David C. Page, Craig C. Mello, Michelle A. Carmell

## Abstract

GCNA proteins are expressed across eukarya in pluripotent cells and have conserved functions in fertility. GCNA homologs Spartan/DVC-1 and Wss1 resolve DNA-protein crosslinks (DPCs), including Topoisomerase-DNA adducts, during DNA replication. We show that GCNA and Topoisomerase 2 (Top2) physically interact and colocalize on condensed chromosomes during mitosis, when Spartan is not present. We show that *C. elegans gcna-1* mutants are sensitive to Top2 poison and accumulate mutations consistent with low fidelity repair of DNA damage, leading to loss of fitness and fertility over generations. We also demonstrate that mouse GCNA interacts with TOP2, and *Gcna*-mutant mice exhibit abnormalities consistent with the inability to process DPCs, including chromatin condensation and crossover defects. Together, our findings provide evidence that GCNA maintains genomic integrity by processing Top2 DPCs in the germline and early embryo, where the genome is challenged with an increased DPC burden.

## Introduction

DNA in all living systems is exposed to damage from both endogenous and exogenous sources. The resulting mutations are particularly consequential in pluripotent cells and germ cells. Mutations in pluripotent cells can contribute to somatic phenotypes such as premature aging, cancer and developmental defects. Mutations in germ cells are acutely harmful as these cells are uniquely tasked with passing their genomes to the next generation, a process critical for both short term reproductive success and long term fitness and survival of a species. Germ cells cope with insults that somatic cells never encounter — hundreds of programmed meiotic double-strand breaks, exquisite chromosome movements, homologous recombination, and massive exchange of histones. As such, specialized pathways have evolved to protect the genomic integrity of pluripotent cells and germ cells (Cerutti and Casas-Mollano, 2006; Juliano et al., 2011; Juliano et al., 2010; Shabalina and Koonin, 2008; van Wolfswinkel, 2014).

We previously discovered the ancient GCNA protein family that is present across eukarya in cells carrying a heritable genome, including pluripotent cells and germ cells of diverse multicellular animals (Carmell et al., 2016). *Gcna* mutations in both *C. elegans* and mice significantly impact reproduction, suggesting that GCNA has functioned in the germline for at least 600 million years (Carmell et al., 2016). GCNA proteins belong to a larger family that includes Spartan (also known as DVC-1) and Wss1, which have been implicated in DNA damage responses through the DNA-protein crosslink (DPC) repair pathway (Carmell et al., 2016; Fielden et al., 2018). DPC repair eliminates proteins that are inappropriately crosslinked to DNA (Barker et al., 2005). Endogenous reactive aldehydes, ionizing radiation, UV light, chemotherapeutics, chemical crosslinkers, and trapped enzymatic intermediates can cause DNA-protein crosslinks (Stingele et al., 2015). DPCs interfere with transcription, unwinding, replication, and repair of DNA, as they cannot be bypassed by DNA tracking enzymes (Nakano et al., 2013; Nakano et al., 2012; Yudkina et al., 2018). The SprT domains of Spartan and Wss1 proteolyze the protein components of DPCs to make way for downstream repair (Balakirev et al., 2015; Centore et al., 2012; Davis et al., 2012; Juhasz et al., 2012; Kim et al., 2013; Machida et al., 2012; Mosbech et al., 2012; Stingele et al., 2014).

Although they process a number of substrates, topoisomerases are major targets of Spartan and Wss1(Lopez-Mosqueda et al., 2016; Maskey et al., 2017; Stingele et al., 2014; Vaz et al., 2016). Topoisomerases modify DNA topology and are necessary for DNA replication, transcription, recombination, chromatin condensation, and chromosome segregation (Wang, 1996). Top1 and Top2 make single and double-stranded DNA breaks, respectively, and have covalent reaction intermediates in which a tyrosine in the active site is crosslinked to DNA (Champoux, 2001; Wang, 2002). Abortive reaction events leave DPCs that must be resolved before they are encountered by DNA tracking enzymes such as helicases or polymerases (Deweese and Osheroff, 2009).

Spartan, which is highly expressed during S phase, is a replication-associated DPC repair protein that travels with the replisome and resolves DPCs blocking replication forks (Morocz et al., 2017; Vaz et al., 2016). Although many DPCs are resolved during S phase, DPCs are also generated in other phases of the cell cycle when Spartan is absent due to degradation by APC-Cdh1 (Mosbech et al., 2012). Top2 DPCs present a likely target outside of S phase for which a protease has heretofore not been identified. Top2 is highly expressed during the G2/M phase of the cell cycle in proliferative cells, where it is necessary for chromatin condensation and proper separation of sister chromatids through the release of helical torsion and resolution of knots and catenanes (DiNardo et al., 1984; Li et al., 2013; Maeshima and Laemmli, 2003; Uemura et al., 1987; Uemura and Tanagida, 1986; Woessner et al., 1991).

Top2 is abundant throughout the germline where it has several germline specific functions including separation of recombined chromosomes, crossover interference, histone exchange, and sperm chromatin condensation (Akematsu et al., 2017; Benkert et al., 2011; Hartsuiker et al., 1998; Hughes and Hawley, 2014; Jaramillo-Lambert et al., 2016; Leduc et al., 2008; Marchetti et al., 2001; Marcon and Boissonneault, 2004; Mengoli et al., 2014; Rathke et al., 2007; Tateno and Kamiguchi, 2001). In accordance with these critical functions, topoisomerase dysfunction during meiosis in a wide array of organisms including yeasts, mammals, fly, and worm causes chromosome segregation defects that result in aneuploidy and chromosome breakage in spores and gametes (Benkert et al., 2011; Hartsuiker et al., 1998; Hughes and Hawley, 2014; Jaramillo-Lambert et al., 2016; Marchetti et al., 2001; Mengoli et al., 2014; Tateno and Kamiguchi, 2001). Top2 also functions in the early embryo where it is necessary for paternal chromatin remodeling and activation of the germline zygotic genome after fertilization (Tang et al., 2017; Wong et al., 2018). In addition to Top2, germ cells also express a specialized topoisomerase, Spo11. Spo11 is responsible for programmed meiotic double strand breaks and sperm chromatin condensation (Akematsu et al., 2017; Keeney et al., 1997). Taken together, early embryo and germline genomes are expected to carry an extra burden of Topoisomerase DPCs relative to cycling somatic cells.

Here we show that in the absence of GCNA, the genome is subject to mutations, including multi-kilobase deletion, inversions, and copy number increases, which cause deterioration of the genome over successive generations. This phenotype is consistent with that of *dvc-1* and points toward a role in DPC repair. Indeed *dvc-1;gcna-1* double mutants exhibit a synthetic sterility phenotype. Herein, we identify Top2 as a major target of GCNA. We show that in *C. elegans* GCNA-1 and TOP-2 physically interact and colocalize on condensed chromosomes during M phase of the cell cycle, at which time DVC-1 is not present due to cell cycle regulation. Consistent with the idea that GCNA is primarily responsible for processing TOP-2 DPCs, *gcna-1* mutants are sensitive to TOP-2, but not TOP-1, poison. Mouse GCNA also interacts with TOP2, and *Gcna*-mutant mice exhibit abnormalities that are consistent with the inability to process DPCs, including aberrant chromatin condensation, persistent DNA damage, crossover anomalies, and sperm chromatin condensation defects. Together, our findings support the model that in the germline and early embryo, GCNA buttresses Spartan/Dvc-1 anti-DPC activity by processing Top2 DPCs.

## Results

### *gcna-1* mutants exhibit a distinct germline phenotype associated with genomic decline

Based on phylogenetic analyses, we previously reported that GCNA proteins are members of a family comprised of Wss1 and Spartan proteases that are implicated in DPC repair (Carmell et al., 2016) (Figure 1A). GCNA, Wss1, and Spartan share highly homologous protease domains and large, rapidly evolving disordered regions (Carmell et al., 2016). In order to investigate whether GCNA and other members of this family have conserved function, we further characterized the phenotype of *C. elegans* strains bearing *gcna-1* mutations. Like most animals, in addition to *gcna-1, C. elegans* has a single additional related gene that is most similar to Spartan (*dvc-1*). While both *gcna-1* and *dvc-1* mutant *C. elegans* display decreased brood sizes, *dvc-1* broods are markedly smaller (Figure 1B)(Carmell et al., 2016; Mosbech et al., 2012). To determine if *gcna-1(ne4356)* mutants display germline morphological defects, *gcna-1(ne4356)* mutants were grown at 20°C and 25°C, dissected and stained with the mitotic proliferative marker, phospho-histone 3 (PH3), and with the P-granule marker, PGL-1 (Brangwynne et al., 2009; Hendzel et al., 1997). P-granules are perinuclear RNA-protein granules that serve as a hub of post-transcriptional germline control (Brangwynne et al., 2009; Voronina, 2013). At both temperatures we observed wild-type distributions of both markers (Figure S1A and S1B), suggesting that the germlines of *gcna-1(ne4356)* mutants exhibit grossly wild-type organization.

**Figure 1:**
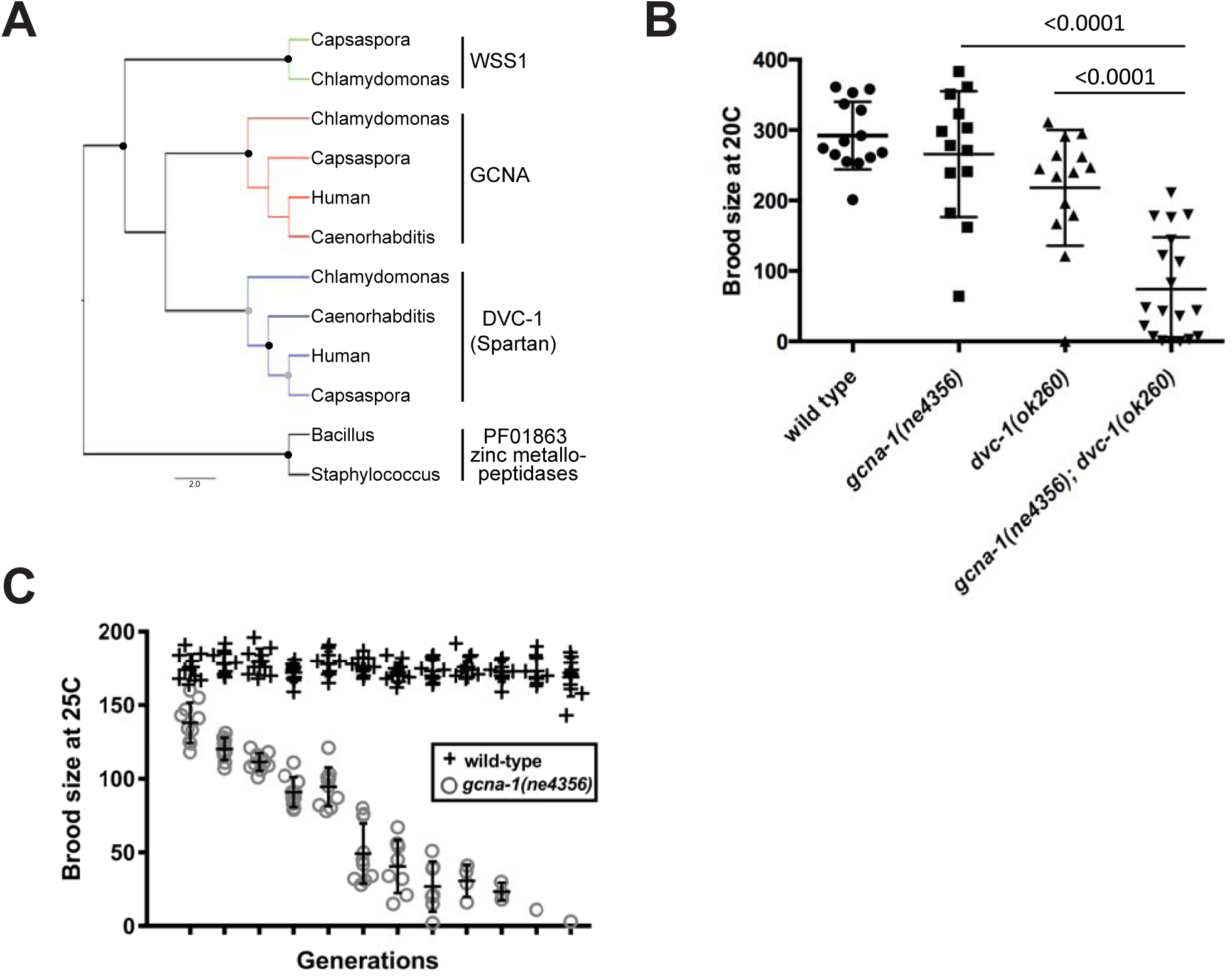
*gcna-1* mutants exhibit a distinct germline phenotype associated with genomic decline. (A) Maximum likelihood phylogenetic tree showing GCNA is closely related to DVC-1 (Spartan) and Wss1 proteases. Gray and black circles indicate bootstrap values higher than 750 and 900 (out of 1000), respectively. Zinc metallopeptidases from the same clan but different family were used as an outgroup. (B) Brood size comparison between *gcna-1(ne4356), dvc-1(ok260)*, and *gcna-1(ne4356);dvc-1(ok260)* double mutants. (C) Brood sizes of wild-type and *gcna-1(ne4356*) mutant worms across twelve generations at 25°C. Each symbol represents the number of progeny derived from a single hermaphrodite. Bars indicate mean +/-standard deviation.

Gradual loss of genomic or epigenetic integrity in germ cells can result in sterility over successive generations, a phenotype termed “germline mortality” (Ahmed and Hodgkin, 2000; Harris et al., 2006; Meier et al., 2006). Interestingly, despite an apparently mild phenotype in early generations, *gcna-1(ne4356)* mutants grown at 25°C have a mortal germline, where brood sizes become progressively smaller and the population fails to survive beyond 12 generations (Figure 1C). To analyze this phenotype further, *gcna-1(ne4356)* mutants were grown at 25°C and subjected to acridine orange staining which selectively stains apoptotic cells in the germline (Gumienny et al., 1999). When compared to wild-type animals, *gcna-1(ne4356)* mutants displayed moderately elevated levels of germ cell apoptosis (Figure S1C). These findings are consistent with a role for GCNA-1 in maintaining germ cell viability and germline immortality.

The GCNA-1 homolog DVC-1 is also expressed in the germline. We therefore wished to investigate whether the two genes have redundant functions. For this analysis, we used *dvc-1(ok260)*, a presumptive null allele, which removes the first three exons as well as portions of the promoter. Consistent with parallel or redundant functions, we found that the fertility defects in *gcna-1;dvc-1* double mutants were significantly more pronounced than in either *gcna-1* or *dvc-1* mutants alone (Figure 1B). Taken together, our results indicate that *gcna-1* and *dvc-1* have partially overlapping functions required for fertility.

### *gcna-1* is required for response to replication stress

The GCNA-1 homolog DVC-1 has been implicated in the response to replication stress caused by both ultraviolet (UV) irradiation and hydroxyurea (HU) (Mosbech et al., 2012). For example, the mouse DVC-1 ortholog (SPRTN) is recruited to sites of UV-induced DNA damage and is necessary for lesion bypass at the resultant stalled replication forks (Centore et al., 2012; Maskey et al., 2014). *C. elegans* DVC-1 also localizes to sites of UV damage, and *dvc-1* mutant worms exhibit increased embryonic lethality compared to wild-type controls in response to UV irradiation (Mosbech et al., 2012). Hydroxyurea inhibits DNA synthesis by depleting dNTPs, which causes arrested replication forks and ultimately increased levels of double-strand breaks (Singh and Xu, 2016). HU treatment also increases the levels of abortive Top2 reaction intermediates covalently bound to DNA (Lee et al., 2012). Upon HU treatment, the human DVC-1 ortholog is recruited to foci containing PCNA at blocked replication forks (Davis et al., 2012; Mosbech et al., 2012). In *C. elegans*, *dvc-1* mutant larvae treated with HU have higher rates of sterility compared to wildtype animals suggesting that *dvc-1* is required to cope with replication stress (Mosbech et al., 2012).

Because *gcna-1* is partially redundant with *dvc-1* for fertility, we sought to directly compare *gcna-1* and *dvc-1* mutants’ response to exogenous DNA damage. Consistent with previous reports, we observed increased sensitivity of *dvc-1* mutants to both UV irradiation and HU treatment (Figure 2A,B, Figure S2). By contrast, hatching rates of *gcna-1* mutant embryos were unaffected by UV irradiation (Figure 2A), while HU treatment increased embryonic lethality compared to wild-type controls (Figure 2B, Figure S2).

**Figure 2:**
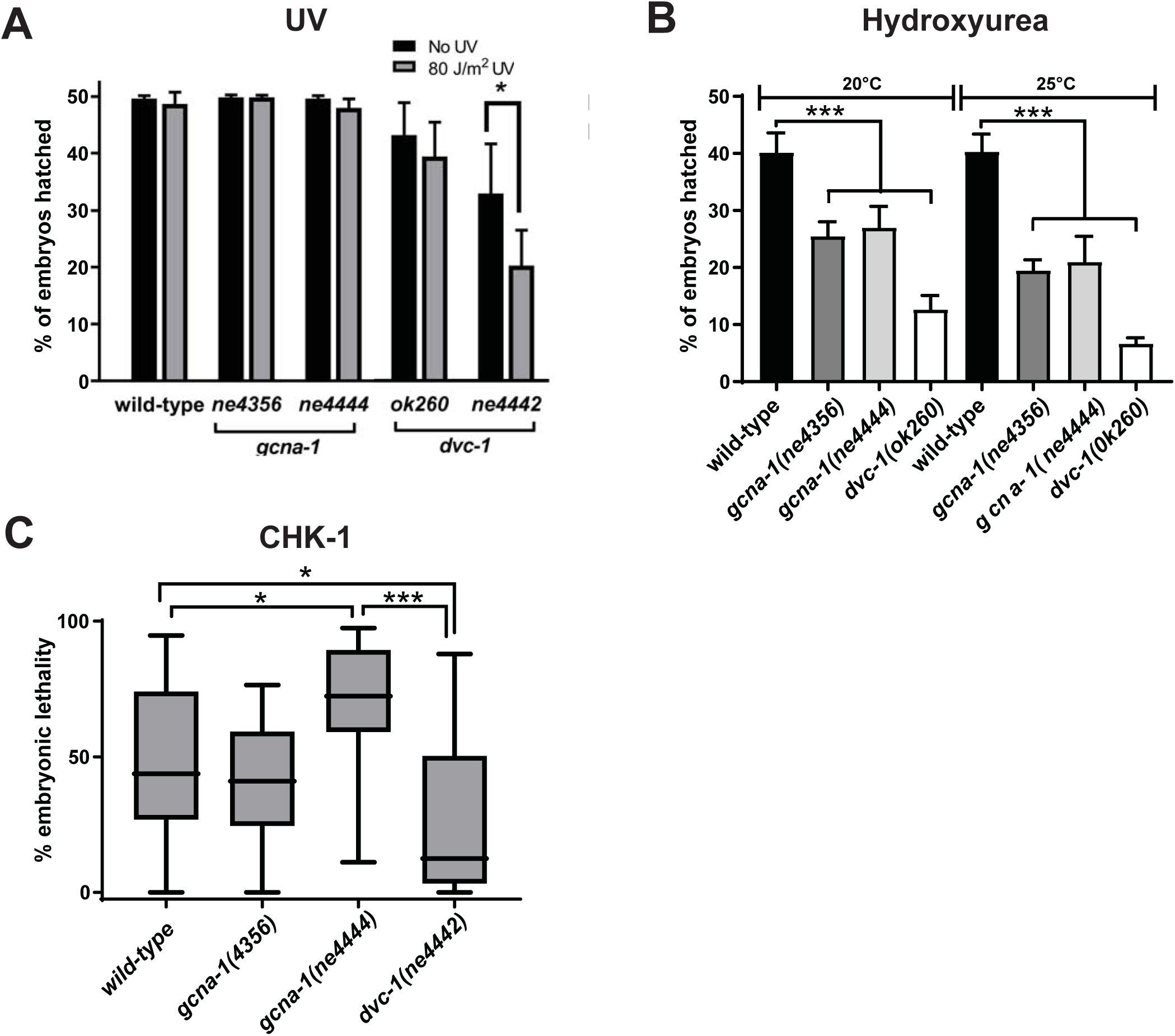
*gcna-1* is required for response to replication stress. (A) Hatching rate of embryos laid by adults exposed as L4 larvae to ultraviolet (UV) irradiation. (B) Hatching rate of embryos laid by adults after 20 hours of exposure to hydroxyurea. Error bars in A and B= SEM. (C) Knockdown of *chk-1* by RNAi in *gcna-1* mutants results in embryonic lethality. Values are normalized to the mean hatching rate of untreated controls. Box depicts 25th and 75th percentiles, and median. Whiskers represent the minimum and maximum values.

Checkpoint kinase, CHK-1, is a critical part of response to HU in *C. elegans* embryos (Brauchle et al., 2003). Upon DNA damage, CHK-1 is activated and prevents entry into mitosis until damage has been repaired. Even in the absence of exogenous insult, *chk-1* is required for successful completion of DNA replication, as knockdown of *chk-1* results in embryonic lethality due to premature entry in to M phase (Kalogeropoulos et al., 2004). Interestingly, SPRTN (DVC-1) has been recently shown to be involved in CHK-1 activation under normal DNA replication conditions (Swagata Halder, 2018). We therefore asked how *dvc-1* and *gcna-1* mutants respond to *chk-1* RNAi-induced depletion under otherwise normal conditions. In agreement with previous reports, we found that knockdown of *chk-1* by RNAi leads to highly penetrant embryonic lethality in wild-type animals (Figure 2C) (Kalogeropoulos et al., 2004). In *dvc-1* mutants, we found a substantial rescue of the embryonic lethality phenotype that is consistent with its proposed role in activating CHK-1. However, in *gcna-1* mutants we did not observe such a rescue and instead observed a slight increase in embryonic lethality. This differential interaction with *chk-1* under normal growth conditions suggests that unlike *dvc-1*, which appears to have a distinct function upstream of *chk-1*, *gcna-1* likely acts downstream of the checkpoint.

### Absence of *gcna-1* causes a potent mutator phenotype

We observed an increased rate of spontaneous mutant phenotypes including abnormal vulva and body morphology during long term culture of *gcna-1* mutant animals (Data not shown). This observation, along with the mortal germline and *him* phenotypes (Carmell et al., 2016), could be explained by an increase in the spontaneous mutation frequency in the *gcna-1* germline. To explore this possibility more directly, we carried out a sensitive genetic assay for measuring spontaneous mutations (Harris et al., 2006). This assay utilizes the semi-dominant gain-of-function *unc-58(e665)* allele, which produces small, paralyzed worms with a shaker phenotype (Figure 3A) (Brenner, 1974; Harris et al., 2006). The paralyzed phenotype of *unc-58(e665)* mutants can be suppressed by either intragenic loss of function or by extragenic mutations that are easily identified as animals with wild-type motility (Hodgkin, 1974). Genetic backgrounds that cause an increase in the number of spontaneous mutations are expected to produce more revertants than would occur in *unc-58(e665)* alone. Consistent with the idea that GCNA-1 promotes genome integrity, we found that spontaneous revertants of *unc-58(e665)* occurred 10.6 and 12 times higher in the *gcna-1(ne4356)* and *gcna-1(ne4444)* mutant backgrounds (respectively), as compared to *unc-58(e665)* alone. We also found that *dvc-1(ok260)* mutants also exhibited an increased rate of *unc-58(e665)* reversion, to 35 times higher than the background rate (Figure 3B).

**Figure 3:**
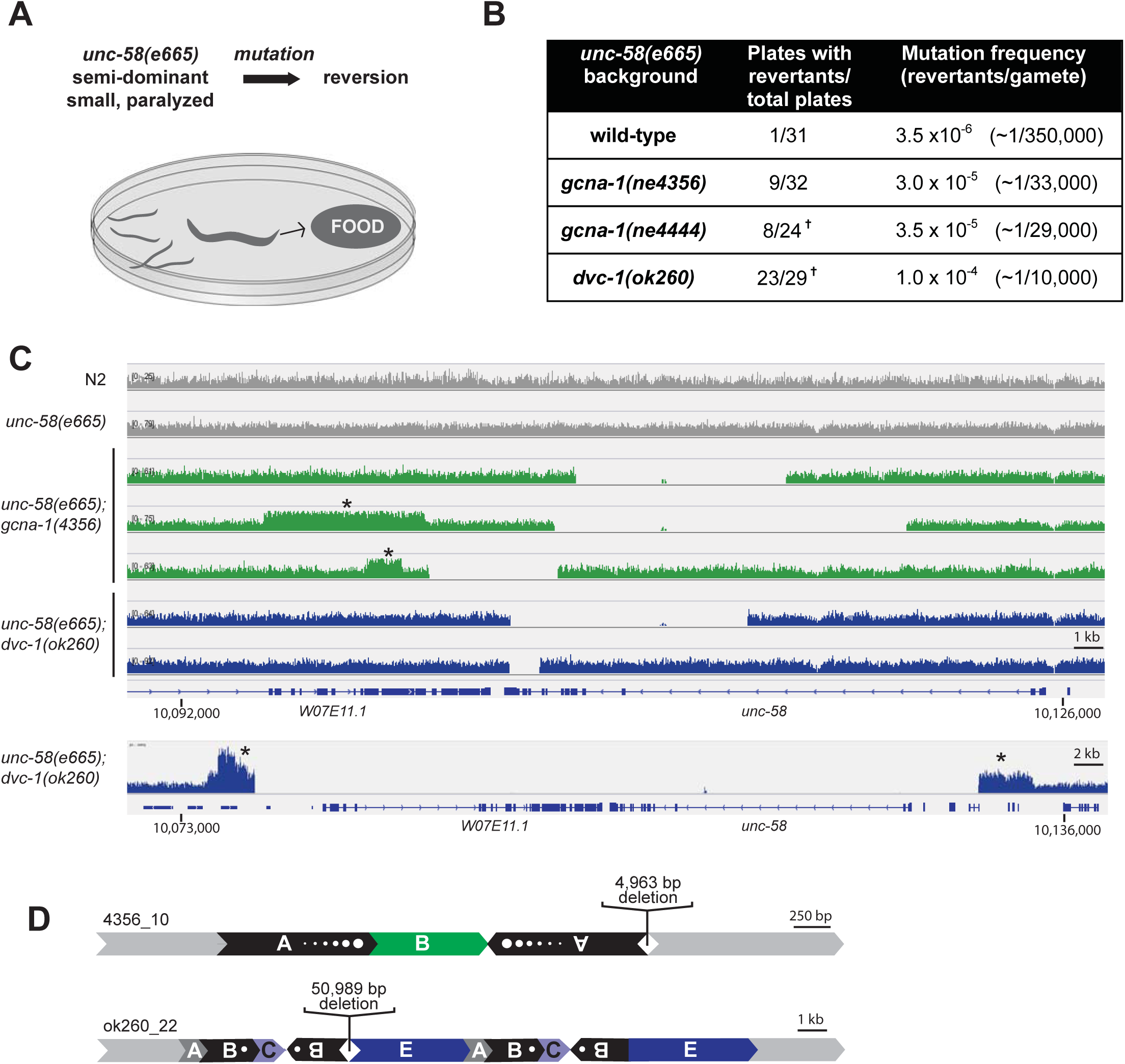
Absence of *gcna-1* causes a potent mutator phenotype. (A) Schematic of *unc-58(e665)* mutator assay. (B) Frequencies of spontaneous mutation as determined in the mutator assay. Crosses denote that at least two independent reversion events took place in a single plate, as evidenced by two distinct reverted phenotypes (N=1 plate for *gcna-1(ne4444)* and N=4 plates for *dvc-1(ok260)*. (C) Whole genome sequencing coverage surrounding the *unc-58* gene. Deletions are indicated by absence of sequencing reads. Asterisks indicate regions with increased copy number relative to the surrounding genome. Mutant samples are (from top to bottom): green, gcna-1 ne4356_5, ne4356_4, ne4356_10; blue, dvc-1 ok260_6, ok260_7, ok260_22. Panel is modified from the Integrative Genomics Viewer (Broad Institute). (D) Structural rearrangements at the *unc-58* locus in revertant lines.

In order to determine the nature of the mutations in the *unc-58* gene, we tiled the gene by PCR with primers at 1kb intervals (Data not shown). We found large, multi-kilobase deletions in some samples but were unable to amplify breakpoints in others. We suspected structural rearrangements and therefore sequenced the entire genome of three *gcna-1(ne4356);unc-58(e665)* revertants and three *dvc-1(ok260);unc-58(e665)* revertants. We aligned read pairs to the *C. elegans* reference genome (ws268; N2 strain) and ran three copy number callers and three structural variant (SV) callers (see Methods, Table S1). We used a consensus approach for SV calls in which candidate rearrangements had to be identified in at least two out of three callers. Paired sequencing reads were considered to be discordantly mapping with respect to the reference sequence if they fell into one of three categories: 1. Inferred insert size was larger than expected based on the average insert size in the sequencing library, indicating a possible deletion or translocation. 2. Both reads mapped to the same strand, implying the existence of an inversion. 3. Reads mapped to opposite strands but in the wrong orientation relative to the reference, implying the presence of a duplication or translocation.

Sequencing read depths in the region surrounding *unc-58* are shown for wild-type (N2), the *unc-58(e665)* parental strain, and the selected revertants (Figure 3C). Deep sequencing confirmed the presence of deletions in the *unc-58* gene of approximately 5-13.5 kb in *gcna-1(ne4356);unc-58(e665)* revertants, and from 1-51 kb in the *dvc-1(ok260);unc-58(e665)* revertants (Table S2). The presence and sizes of these deletions are consistent with those previously found in the *unc-58* locus in worms carrying mutations in DNA damage response genes using the same assay (Harris et al., 2006).

In two *gcna-1* and one *dvc-1* mutant revertants, we observed that blocks of DNA adjacent to the deletions in *unc-58* had more than 1.8X the expected number of sequencing reads, suggesting that *de novo* duplications had occurred adjacent to the deletions. Further analysis revealed the complex nature of these duplications (Figure 3D, Figure S3). In one *gcna-1* mutant revertant (4356_4), an ~7.5 kb block of sequence situated ~5 kb upstream of the breakpoint was duplicated in tandem. In another, (4356_10), an ~1.4 kb block of sequence that normally exists in one copy 1.1 kb upstream of the breakpoint was duplicated and inserted in an inverted orientation immediately adjacent to the breakpoint. The most complex rearrangement in *unc-58* occurred in the *dvc-1* mutant background (ok260_22). Like one *gcna-1* mutant revertant (4356_10), this *dvc-1* revertant has a duplication of a segment upstream of the breakpoint inserted in an inverted orientation adjacent to the breakpoint. The resultant conglomeration of five segments was then duplicated in tandem, resulting in three novel junctions in the genome (Figure 3D, Figure S3).

We wished to determine the extent of the DNA damage that had accumulated over several generations in our revertant lines, so we examined the prevalence of structural variants, including deletions, copy number increases, inversions and translocations across the entire genomes of *gcna-1(ne4356);unc-58(e665)* and *dvc-1(ok260);unc-58(e665)* revertants. We identified one 2255 bp homozygous deletion shared by all three *gcna-1* revertant lines, indicating that it was either pre-existing in the line or derived soon after the cross with *unc-58(e665)*. Likewise, we found 6 homozygous deletions ranging from 574 bp to 13,231 bp shared by all three *dvc-1* mutant lines, as well as one *de novo* 2375 bp deletion unique to ok260_22. Two deletions in the *dvc-1* mutant background are part of complex rearrangements including duplications and inversions similar to those found in *unc-58* (Table S2). Overall, the size and nature of the deletions found *gcna-1(ne4356);unc-58(e665)* revertants was similar to those found in *dvc-1(ok260);unc-58(e665)* revertants.

We also found evidence of duplications, inversions, and translocations in mutant lines that were not present in controls (Table S3). We found nine *de novo* structural variants in *gcna-1(ne4356);unc-58(e665)* revertants, all of which were inversions. The inversions ranged in size from approximately 1 to 7.5 kb. We also found fourteen *de novo* variants in *dvc-1(ok260);unc-58(e665)* revertants that were comprised of ten inversions, two tandem duplications, and two complex rearrangements with signatures of inversion, duplication, and deletion. Inversions in the *dvc-1* mutant background ranged from approximately 800 bp to 8 kb, while one inversion and the tandem duplications were larger (26 kb and 10-12 kb, respectively). We did not find homozygous translocations between chromosomes in any of our mutant strains.

In addition to discordant reads that provided evidence for homozygous structural variants, we found an abundance of rare discordant reads in both *gcna-1* and *dvc-1* mutant lines when compared to wildtype and *unc-58(e665)* backgrounds (Table S4). In the N2 and *unc-58(e665)* control samples, 0.15% and of the uniquely mapping read pairs in each were discordant, while mutant samples had between two and eight times that amount, a difference of hundreds of thousands of discordant reads (Chi-squared analysis; p=0 for all pairwise comparisons). When comparing *gcna-1* and *dvc-1* mutant samples to each other, we found that they were equally enriched in discordant reads. In *gcna-1* mutant samples, an average of 0.51±0.1% of reads were discordant, while an average of 0.81±0.4% of the reads mapped discordantly in *dvc-1* mutant samples.

We also found that the makeup of the discordant read populations was similar across all categories of discordance between *gcna-1* and *dvc-1* mutant lines. Of the discordant reads that mapped to the same chromosome, the average percentage of discordant reads suggesting inversion events was 35±18% and 42±2.3% in *gcna-1* and *dvc-1* mutant lines, respectively. The average percentage suggesting deletions was 29±20% and 13±7.7%, and those suggesting duplications or translocations were 3.9±0.7% and 4.6±0.4% in *gcna-1* and *dvc-1* mutant lines, respectively. We also found that reads suggesting rare interchromosomal translocations were equally common in both mutant lines. Of the total number of discordant reads, an average of 53±18% and 69±12% of discordant reads in *gcna-1* and *dvc-1* mutant backgrounds mapped to two different chromosomes. Overall, these results provide evidence for similar mutational profiles in *gcna-1* and *dvc-1* mutants.

We wished to determine if the discordant reads, both those that led to structural variant calls and those that represented rare rearrangements, were randomly distributed in the genome. We found that discordant reads in our mutant lines often mapped to complex regions of the genome containing multiple copies of genes in the same family, which are inherently unstable (Table S3, S4). Specifically, many of these regions were comprised of palindromes ranging in size from 1-30 kb. Several of the structural variants found in mutant lines appear to result from rearrangements between two inverted copies of the same gene (Table S3). For example, ChrIV: 5845834-5849201 is an approximately 3000 bp inversion between *sams-3* and *sams-4* genes, which are highly homologous and are situated in an inverted orientation relative to each other.

Many of the lesions in *gcna-1* and *dvc-1* mutants are consistent with the error prone process of break induced replication, which is known to routinely result in duplications and fold back inversions (Costantino et al., 2014; Malkova and Haber, 2012). Interestingly, we detected an association between GCNA and both the catalytic subunit of DNA polymerase delta and POLD3, which is dispensable for typical replication but essential for break induced replication, in our mouse ES cell mass spectrometry dataset (Costantino et al., 2014; Lydeard et al., 2007)(Table S5).

### GCNA-1 is cell cycle regulated and localizes to condensed chromosomes during M phase

The mammalian *dvc-1* ortholog, Spartan, is a constitutive component of the replication machinery and is required for replication-coupled DPC repair (Lopez-Mosqueda et al., 2016; Maskey et al., 2017; Morocz et al., 2017; Vaz et al., 2016). Spartan is expressed primarily during the S and G2 phase of the cell cycle; it is regulated by APC-Cdh1 and degraded in mitosis (Mosbech et al., 2012). We carried out cell cycle analysis of GCNA expression in mouse embryonic stem cells and found that GCNA protein is also cell cycle regulated. GCNA expression is low in G1 and enriched in S and G2/M phases of the cell cycle (Figure 4A). In *Schizosaccharomyces pombe*, studies in synchronized cells provide finer cell cycle resolution for GCNA and the other pombe SprT protein, Wss1. While *wss1* transcripts rise during S and reach their highest level in G2, GCNA (*SPBC19G7.04*) has a clear peak in expression during M phase (Figure 4B)(Bahler, 2005).

**Figure 4:**
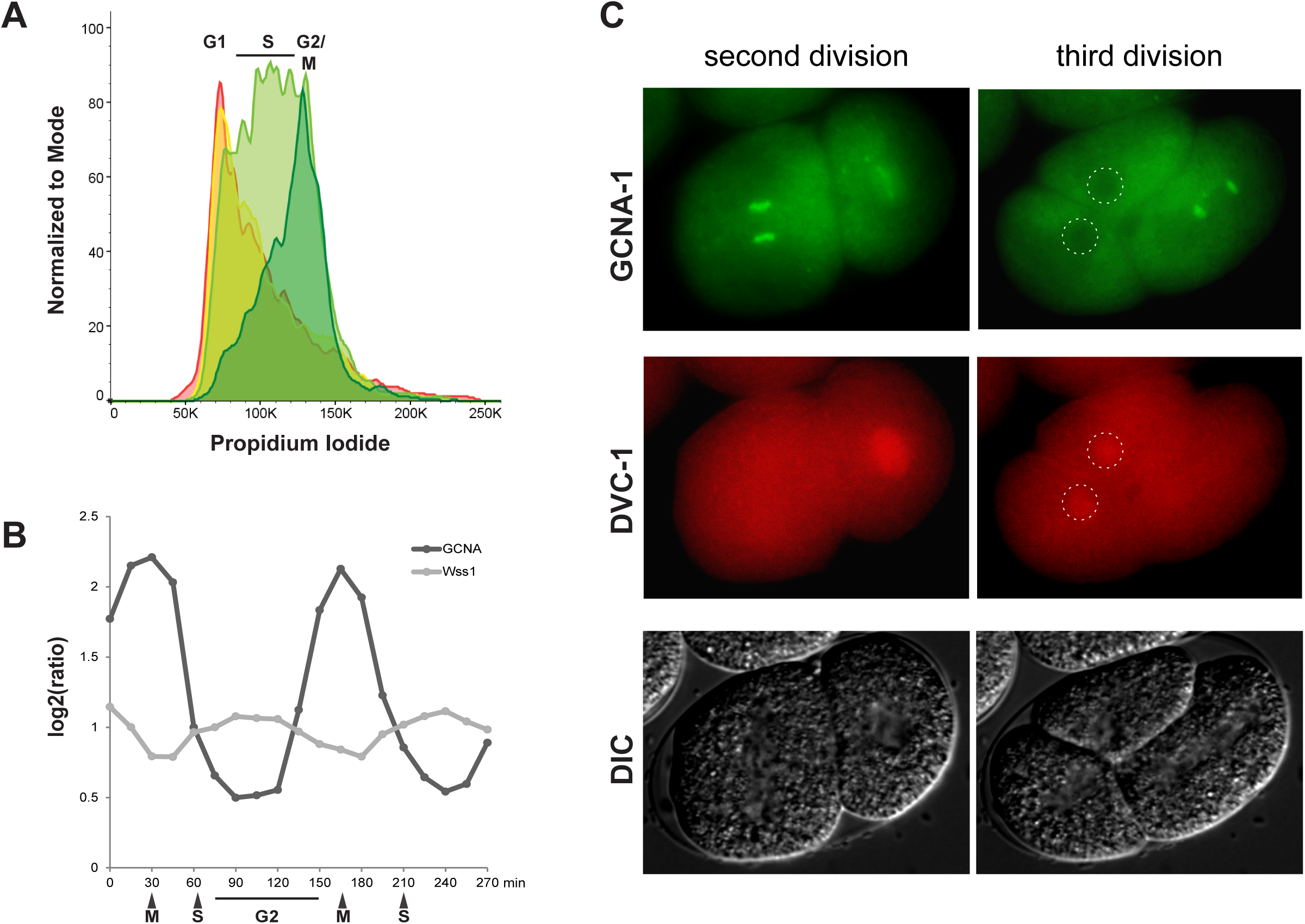
GCNA is cell-cycle regulated and localizes to condensed chromosomes during M phase. (A) GCNA protein expression in mouse embryonic stem cells. Colors correspond to relative GCNA protein level as measured by immunostaining with anti-GCNA antibody followed by flow cytometry and are defined as follows: red (no GCNA expression), yellow (low), light green (medium), dark green (high). (B) Cell-cycle regulated expression of *Schizosaccharomyces pombe* GCNA and Wss1 transcripts shown after synchronization by Cdc25 block and release (Rustici et al., 2004). The timing of mitosis (M), S, and G2 phases are indicated. (C) Localization of GFP::GCNA-1 and DVC-1::mCherry in live *C. elegans* embryos during the second and third cell divisions. Nuclei in ABa and ABp cells are indicated by dashed circles.

To investigate the possibility of a cell-cycle regulated hand off between Spartan and GCNA in *C. elegans*, we set out to explore the localization of DVC-1 and GCNA-1 using fluorescently tagged CRISPR alleles of both proteins (Dokshin et al., 2018). In lines carrying both GFP::GCNA-1 and mCherry::DVC-1 alleles, the two proteins had complementary localization dynamics. Specifically, when mCherry::DVC-1 was clearly enriched in the nucleus, GFP::GCNA-1 was clearly excluded. Upon nuclear envelope breakdown, as mCherry::DVC-1 faded, GFP::GCNA-1 became enriched on the condensed chromosomes, and decorated them through completion of mitosis. After mitosis, GFP::GCNA-1 was once again excluded from the DNA, and replaced by nuclear mCherry::DVC-1 (Figure 4C, Supplementary movie 1). We propose that M phase expression and chromosomal localization may be a conserved feature that distinguishes GCNA from other SprT family members and underlies its role in DPC repair during this phase of the cell cycle.

### Topoisomerase II co-localizes and interacts with GCNA

Spartan and Wss1 are required for processing DPCs consisting of Topoisomerase I reaction intermediates crosslinked to DNA (Maskey et al., 2017; Stingele et al., 2014; Vaz et al., 2016). We carried out immunoprecipitation of the GCNA protein from UV-irradiated mouse embryonic stem cells followed by mass spectrometry and identified TOP2 as a major interactor of GCNA (Figure 5A). Vertebrates encode two Top2 isozymes termed alpha and beta. TOP2 alpha functions in chromosome condensation and segregation like the single TOP-2 in *C. elegans* and yeasts (Austin and Marsh, 1998). The majority of our peptides were derived from TOP2 alpha, but we also recovered peptides from TOP2 beta and TOP1 (Figure 5A, Table S5). Although UV irradiation may have enhanced the interaction between GCNA and topoisomerases, *Gcna*-mutant embryonic stem cells are not sensitized to UV, suggesting that GCNA is not required to process UV-induced lesions (Figure S4).

**Figure 5:**
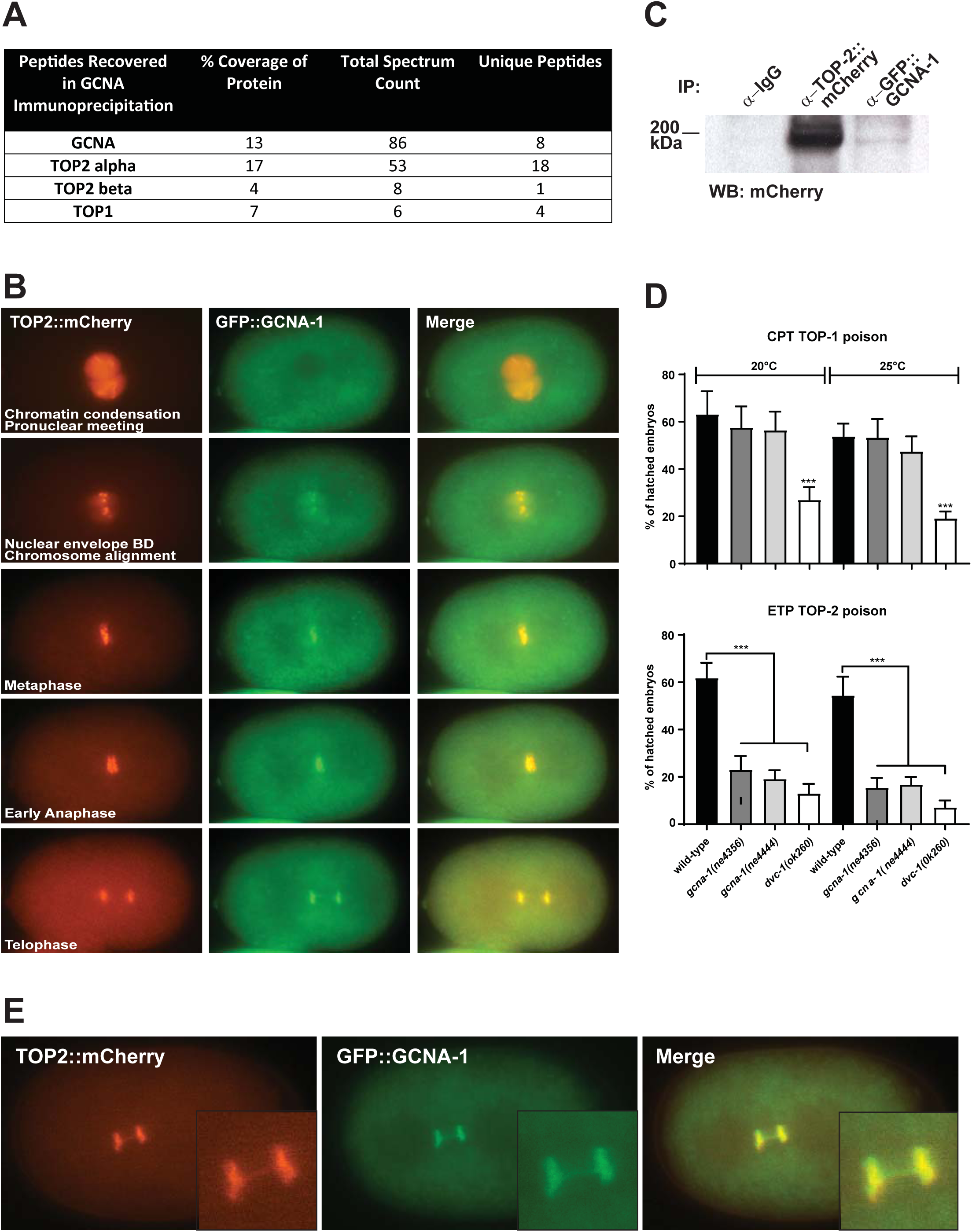
GCNA-1 and TOP-2 physically interact, colocalize on condensed chromosomes, and have a functional relationship. (A) Peptides recovered from anti-GCNA immunoprecipitation from mouse embryonic stem cells followed by mass spectrometry. (B) Live cell imaging of the first embryonic cell ivision in *C. elegans* showing co-localization of TOP-2:mCherry and GFP::GCNA-1. (C). Co-immunoprecipitation of TOP-2::mCherry with GFP::GCNA-1. Immunoprecipitation from *C. elegans* lysates was carried out with anti-GFP antibody and resultant complexes were western blotted with anti-mCherry antibody. (D) Wildtype, *gcna-1* and *dvc-1* mutant L4 larvae were treated with topoisomerase I inhibitor (camptothecin, CPT) and topoisomerase II inhibitor (etoposide, ETP) for 20 hours, and embryo lethality was assayed. (E) Live cell imaging of *C. elegans* embryo showing co-localization of TOP-2:mCherry and GFP::GCNA-1 on anaphase chromatin bridges in the TOP-2::mCherry hypomorphic allele.

Interestingly, in yeast, worm, and mammals, Top2 expression peaks in G2/M and the protein localizes along condensed mitotic chromosomes, as well as along the meiotic prophase axes (Gomez et al., 2014; Jaramillo-Lambert et al., 2016; Kleckner et al., 2013; Maeshima and Laemmli, 2003; Moens and Earnshaw, 1989). Thus, Top2 expression and GCNA expression exhibit similar cell-cycle dependencies. We therefore carried out live cell imaging with GFP::GCNA-1 and TOP-2::mCherry in *C. elegans* and confirmed colocalization of GCNA-1 and TOP-2 on condensed chromosomes during embryonic cell divisions (Figure 5B). In order to confirm the physical interaction between GCNA-1 and TOP-2 suggested by mass spectrometry and colocalization, we conducted co-immunoprecipitation experiments and were able to detect TOP-2::mCherry in complexes isolated from GFP::GCNA-1 immunoprecipitation by western blotting (Figure 5C).

### GCNA-1 mutants are sensitive to TOP-2 but not TOP-1 inhibition

We sought to confirm whether the spatio-temporal co-localization and physical interaction of GCNA and TOP2 reflects the fact that TOP2 DPCs are targets of GCNA during DPC repair. In order to do this, we treated worms with topoisomerase poisons to induce DPCs consisting of trapped reaction intermediates. We investigated both TOP-1 and TOP-2 since we isolated a few TOP1 peptides in our mass spectrometry experiment. We treated worms with camptothecin, a TOP-1 inhibitor, and used embryo hatching rate as a readout of unrepaired DNA damage. We found that *dvc-1(ok260)* worms were more sensitive to camptothecin than wildtype worms, while *gcna-1(ne4356)* and *gcna-1*(*ne4444)* worms were unaffected (Figure 5D). In contrast, treatment with the TOP-2 inhibitor etoposide revealed that mutants in both genes were sensitive to the poison (Figure 5D). Consistent with our protein interaction data from mouse embryonic stem cells, this suggests that DPCs consisting of TOP-2, and not TOP-1, are the primary target of GCNA-1 in *C. elegans*.

Of note, our *C. elegans* TOP-2::mCherry fusion appears to be a hypomorphic allele, as we observed chromatin bridges during mitosis in the early embryo in the TOP-2::mCherry line but never in wildtype (Figure 5E). Top2 is required for decatenation of replicated chromosomes before division and proper separation of sister chromatids in mitosis (DiNardo et al., 1984; Li et al., 2013; Maeshima and Laemmli, 2003; Uemura et al., 1987; Uemura and Tanagida, 1986; Woessner et al., 1991). Temperature sensitive alleles of Top2 or chemical inhibition result in the formation of anaphase chromatin bridges (Cimini et al., 1997; Uemura et al., 1987). Interestingly, we found that GFP::GCNA-2 and TOP-2::mCherry remain on the entangled DNA of bridges in this TOP-2::mCherry hypomorph (Figure 5E, Supplementary movie 2), further suggesting that GCNA and TOP2 have a functional relationship.

### GCNA mutant mouse spermatocyte and spermatid defects are consistent with impaired TOP2 function

In light of the connection we have drawn between GCNA-1 and TOP-2 in *C. elegans*, we examined our previously described *Gcna*-mutant mice for phenotypes consistent with defects in the removal of DPCs (Carmell et al., 2016). Unlike most GCNA orthologs across eukarya, including *C. elegans*, mouse GCNA protein is predicted to be entirely disordered and lacks the protease domain, zinc finger, and HMG box common in other family members. Nonetheless, male mice carrying a mutant GCNA allele are sterile, indicating that significant function lies in the disordered region. Examination of the phenotypes of *Gcna*-mutant mice provided us with the opportunity to examine the function of the disordered region in isolation. Mouse *Gcna* is expressed throughout germ cell development, including when all key events of meiosis are taking place, including chromosome condensation, double strand break formation and repair, synapsis, crossing over, and remodeling of chromatin for packaging into sperm heads (Enders and May, 1994).

Top2 is involved in relieving helical torsion caused by the transcription machinery and is required for efficient transcription (Mondal and Parvin, 2001). Therefore, we examined mRNA populations in wildtype and *Gcna*-mutant mouse testes at two stages during development, postnatal day 8 (p8) when the testes contain mitotic spermatogonia and meiotic cells in the leptotene phase of Meiosis I, and postnatal day 18 (p18), when all stages of meiotic cells are present (Kluin et al., 1982). We found essentially no changes in gene expression between mutant and wildtype; only 33 genes differed in *Gcna*-mutant compared to wildtype at p8, and 80 genes at p18. Transposon expression also did not change (Table S6).

As Top2 is well known for its role in facilitating chromatin condensation, we next examined *Gcna*-mutant spermatocytes for chromatin abnormalities. Top2 protein is abundant in spermatocytes during the leptotene phase of meiotic prophase when chromosomes are actively condensing in preparation for synapsis (Leduc et al., 2008). Mouse seminiferous tubules are organized such that specific cell types always appear together in a section of the tubule, allowing for precise staging of cell types. We first examined histological sections containing leptotene spermatocytes for chromatin abnormalities, and found that *Gcna*-mutant spermatocytes exhibit dramatic premature chromatin condensation when compared to wildtype controls (Figure 6A and Figure S5A). Premature condensation begins in the leptotene stage, by which time the chromatin in mutant germ cells is largely detached from the nuclear membrane and occupying only a small fraction of the nucleus. *Gcna*-mutant chromatin is more compact at leptotene than wildtype chromatin is at the subsequent stage of zygotene, when chromosomes are more condensed and synapsis has begun (Figure S5A). Remarkably, despite these dramatic defects earlier in meiosis, by the pachytene stage, *Gcna*-mutant nuclei recover a nearly wild-type appearance.

**Figure 6:**
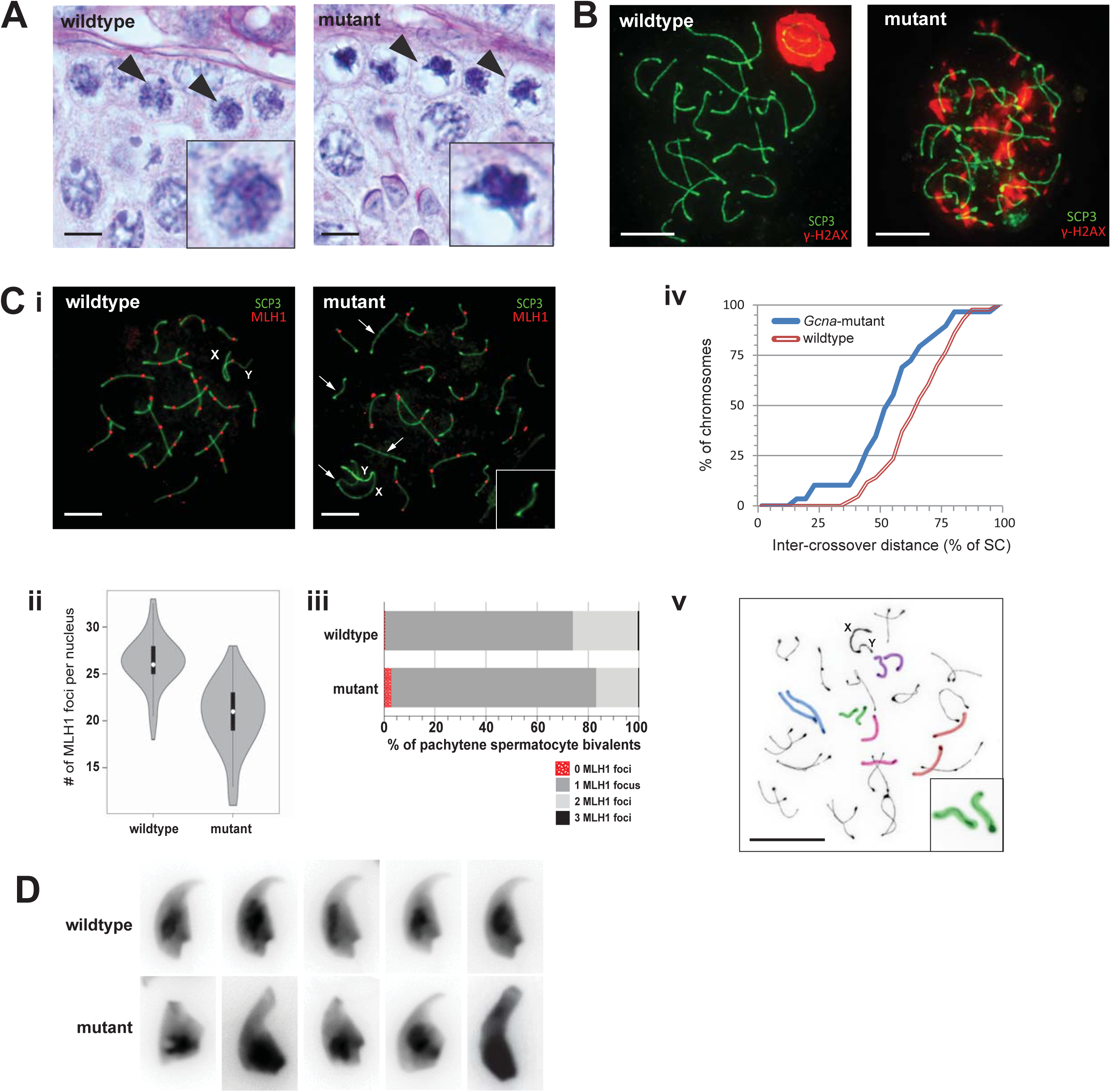
*Gcna*-mutant spermatocytes exhibit DNA damage, crossover defects, and chromatin condensation abnormalities. (A) Hematoxylin and eosin stained testis sections showing Stage IX seminiferous tubules in wildtype (*Gcna+/Y*) and Gcna-mutant (*GcnaDeltaEx4/Y*) animals. Representative leptotene spermatocytes are indicated by black arrowheads and detailed in insets. (B) Pachytene spermatocytes immunostained with synaptonemal complex component SCP3 (green) and DNA damage marker γ-H2AX (red). (C) i. Pachytene spermatocytes stained with SCP3 (green) and recombination nodule marker MLH1 (red). Bivalents lacking MLH1 foci are indicated by white arrows. Inset contains bivalent without an MLH1 focus. ii. Quantification of MLH1 foci in wildtype and *Gcna*-mutant nuclei (Wilcoxan rank sum; p<5e-15). iii. Percentage of pachytene bivalents with 0, 1, 2, or 3 MLH1 foci. iv. Cumulative distribution curves of the distance between two MLH1 foci on Chromosomes 14-19 in wildtype and mutant bivalents normalized to the length of the synaptonemal complex (SC). Crossover interference gamma shape parameter for wildtype is 11.274 and mutant is 8.498 (Two-sided Mann-Whitney; p=0.0051). v. *Gcna*-mutant diplotene spermatocyte stained with SCP3 (black). Univalent chromosome pairs are indicated by colored highlights. (D) Morphology of DAPI-stained wildtype and *Gcna*-mutant sperm heads. (Scale bars, 5 microns)

In light of the genomic instability observed in *C. elegans gcna-1* mutants, we examined meiotic spermatocytes of *Gcna*-mutant mice for hallmarks of DNA damage that would be consistent with aberrant DPC Repair. In wildtype leptotene and zygotene spermatocytes, the DNA damage markers gamma-H2AX, BRCA1, and ATR are found throughout the nucleus due to the presence of programmed meiotic double strand breaks. By the pachytene stage of Prophase I, synapsis is complete, double strand breaks have been resolved, and these proteins no longer occupy bulk chromatin. All three markers subsequently become enriched in the XY body, a specialized chromatin domain containing the mostly asynapsed sex chromosomes (Burgoyne et al., 2007). In order to monitor the progress of meiotic prophase in *Gcna*-mutant spermatocytes, we immunostained spermatocyte spreads with an antibody recognizing SCP3, a component of the synaptonemal complex. We also immunostained for gamma-H2AX, BRCA1, and ATR in order to detect DNA damage and asynapsed chromosomes. Surprisingly, despite the dramatic chromatin condensation in leptotene, ninety percent of *Gcna*-mutant nuclei exhibit normal synapsis and DNA damage resolution. The remainder of spermatocytes exhibit mild asynapsis of one or a few chromosomes, as detected by BRCA1 and ATR staining (Figure S5B). A small number of spermatocytes in *Gcna*-mutant mice retain gamma-H2AX and ATR proteins throughout the nucleus during pachytene even in areas of the nucleus where synapsis has proceeded normally, indicating widespread DNA damage (Figure 6B, Figure S5B).

Crossover interference is the phenomenon by which having a crossover in one spot on a chromosome decreases the probability of another nearby (Hillers, 2004). Top2 has been implicated in crossover interference in yeast (Kleckner et al., 2013). MLH1 is a DNA mismatch repair protein required for crossover formation. MLH1 accumulates in discrete foci along chromosomal axes during pachytene, marking the location of the majority of crossovers (Hassold et al., 2000). In order to assess crossover formation in *Gcna*-mutant mice, we immunostained spermatocyte spreads for MLH1. We found that *Gcna*-mutant mice had significantly fewer MLH1 foci per nucleus, from an average of 26 in wildtype to 21 in mutants (Wilcoxan rank sum; p<5e-15)(Figure 6C.ii). Accordingly, the number of bivalents with no MLH1 focus increased in the mutant relative to wildtype, and the number of bivalents with two foci decreased (Figure 6C.iii).

In order to assess crossover interference, we measured inter-MLH1 focus distance on chromosomes that had more than one MLH1 focus, and graphed the cumulative distribution curves of the distance as a percentage of the length of the entire synaptonemal complex. Mouse chromosomes of different lengths have characteristic differences in crossover distributions, where mean inter-chiasma distances are lower in shorter chromosomes (Hulten et al., 1995). Therefore, we binned chromosomes by length. We found the most significant differences in crossover distributions between wildtype and mutant in the shortest chromosomes (Chr. 14-19)(Figure 6C.iv). We found that crossovers in the *Gcna*-mutant were closer together than wildtype across the entire spectrum of inter-focus distances on these chromosomes. To quantify the degree of crossover interference, we used maximum likelihood fitting of inter-crossover distances to the gamma-distribution and calculated the gamma shape parameter, where a shape parameter of 1 indicates no interference and higher values indicate stronger interference (McPeek and Speed, 1995). We found that wildtype bivalents had a gamma shape parameter of 11.274, while in *Gcna*-mutants it was 8.498, indicating that mutant bivalents exhibit significantly less crossover interference (Two-sided Mann-Whitney; p=0.0051). We found similar results for Chromosomes 11-13, where the wildtype gamma shape parameter was 11.037 and the mutant was 9.479 (Two-sided Mann-Whitney; p=0.0074)(Figure S6) but not for longer chromosomes.

Like mutations in *Gcna*, hypomorphic mutations in mouse *Mre11* and *Nbs1*, components of the MRN complex, reveal defects in double strand break repair, synaptonemal complex integrity, and crossover formation (Cherry et al., 2007). The MRN complex, consisting of MRE11, RAD50, and NBS1, in addition to a number of other roles in double strand break response, is responsible for removal of DPCs that arise from topoisomerase poisons, including TOP1, TOP2, and SPO11 DPCs (Borde, 2007; Deshpande et al., 2016; Hoa et al., 2016; Keeney et al., 1997; Malik and Nitiss, 2004; Sacho and Maizels, 2011). Of note, the *Gcna*-mutant phenotype is highly similar to that of *Zip4h (Tex11*) mutants. ZIP4H is a component of the ZMM complex that interacts with NBS1, forming an association between ZMM and MRN complexes in chromosomal foci (Adelman and Petrini, 2008). Interestingly, we detected an association between GCNA and both RAD50 and the active form of MRE11 in our UV-irradiated mouse ES cell mass spectrometry dataset (Table S5). Of note, we recovered arginine dimethylated MRE11, which is the active form localized to break sites (Boisvert et al., 2005).

DPCs created by both TOP1 and TOP2 poisons are mutagenic to germ cells (Attia et al., 2013; Marchetti et al., 2001). Treatment of mice with etoposide produces DPCs that lead to both structural and numerical chromosome aberrations, including hypo-and hyper-haploid spermatocytes after the first meiotic division (Attia et al., 2002; Marchetti et al., 2001; Marchetti et al., 2006). Aneuploidy produced by etoposide, like the phenotype of *Gcna*-mutant mice, has been associated with effects on recombination (Russell et al., 2000; Russell et al., 2004). Therefore, we examined the integrity of chromosomes in *Gcna* mutants before metaphase I. Pairs of homologous chromosomes are held together during the diplotene phase of meiosis I by chiasmata, physical connections formed by crossovers. Chiasmata are essential for attachment to, and migration towards, opposite spindle poles during metaphase I. As a natural consequence of lacking crossovers, it is expected that bivalents would separate into univalents during diplotene, before metaphase I, leading to missegregation of homologs.

Indeed, we examined at least 50 diplotene spreads of each genotype and found that most *Gcna*-mutant spermatocytes presented with univalents (Figure 6C.v). Overall, 79% of mutant nuclei had at least one achiasmic chromosome compared to 32.5% of wildtype nuclei (Fisher’s Exact Test; p=4e-6). Achiasmic mutant spermatocytes had an average of 4 achiasmic chromosomes per nucleus, compared to 1.8 in wildtype (t-Test; p<0.0005). Of the mutant nuclei lacking at least one chiasma, none lacked chiasma only on the sex chromosomes, 36% had achiasmic autosomes, and 64% had both achiasmic autosomes and sex chromosomes; wildtype achiasmic nuclei had 21%, 43%, and 36% in each category, respectively. These results indicate that the reduced number of crossovers causes *Gcna*-mutant spermatocytes to progress to metaphase I with achiasmic chromosomes. Nondisjunction at metaphase I would consequently lead to aneuploidy in gametes and likely contributes to the sterility of *Gcna*-mutant males.

Top2 is abundantly expressed in elongating spermatids when DNA is undergoing dramatic condensation for packaging into sperm heads (Leduc et al., 2008). In mouse, more than 90% of histones are replaced by transition proteins and then by protamines, ultimately condensing the DNA six times more than in a mitotic chromosome (Jung et al., 2017; Ward and Coffey, 1991). Sperm heads in which chromatin hypercondensation has been disturbed are dramatically misshapen (Gou et al., 2017; Yuen et al., 2014). Topoisomerases, including Top2 and Spo11, create the double strand DNA breaks that facilitate this dramatic germ-cell specific chromatin compaction (Akematsu et al., 2017; Leduc et al., 2008; Marcon and Boissonneault, 2004; Rathke et al., 2007). Top1 may also play a role, as DPCs produced by Top1 poisons induce sperm head abnormalities (R.S et al., 2016). In order to determine whether *Gcna*-mutant sperm have characteristics of topoisomerase dysfunction, we examined sperm isolated from the epididymis of wildtype and *Gcna*-mutant mice and compared their morphology after DAPI staining. *Gcna*-mutant mice generally have fewer sperm in the epididymis, and those that were recovered have an array of abnormal head shapes (Figure 6D). These observations are consistent with failure to execute proper sperm DNA topological rearrangements necessary for full compaction.

## Discussion

The phylogenetic conservation and expression of GCNA proteins suggest an integral role in germ cells and multipotent cells, including those that give rise to germ cells, throughout eukarya (Carmell et al., 2016). Herein we propose that GCNA proteins are required for processing Top2 DNA-protein crosslinks. Typically, cycling cells have a DPC burden of several thousand DPCs per cell even in the absence of exogenous insults (Oleinick et al., 1987). As topoisomerases are the most abundant chromatin-associated proteins after histones, a significant portion of DPCs consist of trapped topoisomerase reaction intermediates (Oleinick et al., 1987; Roca, 2009). Topoisomerases play critical roles in DNA replication as well as in chromosome condensation, decatenation, and segregation and have specialized roles in germ cells. Considering the increased requirement for topoisomerases in germ cells and embryos, it is reasonable to expect that these cells would carry an increased DPC burden, and that specialized pathways may have evolved to process them.

We propose a model wherein Spartan and GCNA complement each other to address an increased DPC burden in the germline and early embryo. While Spartan is primarily active during DNA replication, GCNA acts during mitosis, to ensure robust resolution of DPCs prior to completion of the cell cycle. Our “hand off” model was initially motivated by the partially overlapping phenotypes between *gcna-1* and *dvc-1* mutants, as well as synthetic sterility phenotype of *dvc-1* and *gcna-1* in *C. elegans*. It was further supported by the complementary expression patterns of the two gene products in mice and in yeast, their mutually exclusive and complementary localization pattern in the *C. elegans* embryo, and genetically by the differential interaction of *gcna-1 and dvc-1* with the *chk-1* DNA damage checkpoint. Finally, while Spartan in necessary to resolve Top1 and Top2 DPCs, through chemical biology and biochemical analysis, we were able to elucidate a specific role for GCNA in resolving Top2 DPCs in *C. elegans*, specifically during G2/M. In addition, the meiotic phenotypes of *Gcna*-mutant mice, including persistent DNA damage, decreased crossovers and crossover interference, and chromatin condensation defects echo those produced by both chemical and genetic alterations that cause buildup of DPCs, supporting a role for GCNA in removal of DPCs in the mouse germline. Finally, our bioinformatic analysis has revealed a multitude of genomic alterations in *gcna-1* mutant *C. elegans* with signatures of low fidelity repair of widespread DNA damage that is consistent with buildup of unrepaired DPCs.

We propose that GCNA may be a germ-cell enriched cofactor of typical somatic DPC repair pathways. Topoisomerase adducts are processed through two main avenues: endonuclease digestion and proteolysis followed by reversal of the crosslink. Endonucleases such as MRE11 and CtIP cleave DNA, removing 5’ bulky adducts and leaving 3’ overhangs for downstream repair (Aparicio et al., 2016; Hoa et al., 2016; Neale et al., 2005). Alternatively, tyrosyl-DNA phosphodiesterase 2 (TDP1 and 2) enzymes reverse the crosslink between the tyrosine residue in Top2 and DNA (Cortes Ledesma et al., 2009). However, TDPs can only access the DNA after the majority of the bulky protein adduct has been proteolyzed by upstream proteases, including SprT proteins like Spartan, Wss1, and, likely, GCNA (Gao et al., 2014; Schellenberg et al., 2012).

We propose that most typical GCNA proteins across eukarya may be functioning at DPC sites much like Spartan and Wss1. However, mouse GCNA, which lacks a protease domain, presents a quandary, as it nevertheless retains significant function and appears to be involved in DPC processing in the germline. Importantly, mouse GCNA retains motifs for SUMO interaction (Carmell et al., 2016). SUMOylation and ubiquitylation are common post-translational modifications that regulate recruitment, activity, and stability of damage response proteins at the site of DNA damage (Morris and Garvin, 2017). Both topoisomerases I and II are SUMOylated (Liao et al., 2018; Mao et al., 2000). SUMOylation is necessary for Top2 accumulation along chromosome axes and at centromeres, and defects in SUMOylation result in failure of chromosome segregation (Bachant et al., 2002; Takahashi et al., 2006). SUMOylation of Top2 is also induced in response to topoisomerase-trapping drugs, and has been suggested to be a signal recognized by Top2-responsive checkpoints (Agostinho et al., 2008; Schellenberg et al., 2017). We propose that the SUMO interacting motifs in GCNA mediate interactions with both trapped topoisomerases and with other repair machinery. Given the interaction that we detect between GCNA and two components of the MRN complex, MRE11 and RAD50, we propose that mouse GCNA may be acting as a scaffold to recruit the MRN complex, which is also SUMOylated, to process DPCs (Sohn and Hearing, 2012).

Based on the mutational signature in *gcna-1* mutants, and our protein interaction data connecting GCNA to POLD3, a subunit of polymerase delta that is uniquely required for break induced replication (BIR), we propose that repair downstream of GCNA occurs at least partially through this process. BIR is the process by which the cell re-starts replication from a one-ended double stranded DNA break such as occurs when a replication fork collapses (Malkova and Ira, 2013). Break induced replication is highly error prone and often results in duplications and inversions (Deem et al., 2011), which account for many of the rearrangements in *gcna-1* and *dvc-1* mutants. Consistent with our proposal that *gcna-1* mutation causes buildup of DPCs, break induced replication is involved in repair of DPCs caused by Top1 inhibition (Payen et al., 2008). Mutations in both *gcna-1* and *dvc-1* backgrounds are preferentially located in complex regions of the genome containing palindromes and multi-copy genes. This enrichment is likely not due to a specific function for GCNA-1 and DVC-1 at these sites. Rather, it can be explained by generally poor outcomes of DNA repair pathways in these regions. When breaks occur where multiple homologous regions are in close proximity, during homology searching, the odds of finding the correct copy, in the correct orientation, are relatively low. Some forms of BIR are in fact dependent on the MRN complex, and can lead to recombination between distant inverted repeats, leading to translocations and chromosome fusions (VanHulle et al., 2007). Events such as these may be represented in our hundreds of thousands of discordant sequencing reads, which likely represent genuine, but rare, events that occurred on a single chromosome in a single cell of a single worm. A cell carrying such a fusion would likely be eliminated in the germline due to disruption of meiotic pairing and, if it survived, its genome would likely not be compatible with embryonic development.

We have shown that the germlines of *C. elegans* amass structural damage across generations, and that mutant mice are prone to chromosome nondisjunction, raising the possibility that *Gcna* deficiency could underlie mutations in humans. Mutations in the Spartan gene in humans cause Ruijs-Aalfs syndrome, which is associated with premature aging and liver cancer due to DNA damage and chromosomal instability (Lopez-Mosqueda et al., 2016; Maskey et al., 2014). We have shown here that *Gcna* and Spartan mutations cause similar genomic instability. Because *Gcna* is primarily expressed in the germline while Spartan is expressed more ubiquitously in humans, *Gcna* mutants are more likely to have germline than somatic phenotypes. Human *Gcna* is on the X chromosome, and, based on our insights from the mouse and worm, one would expect that males carrying hypomorphic or null *Gcna* alleles would have compromised fertility, and even sterility. However, offspring arising from *Gcna*-mutant damaged germline genomes might resemble Spartan mutants with regard to somatic phenotypes, as children could have significant genome alterations that originated in the germline of their father. Taken together, our results suggest that GCNA proteins are critical across a wide range of eukaryotic species for ensuring both short term reproductive success and long term fitness and survival of species.

## Supporting information

Table S1 Alignment Metrics

Table S2 Homozygous Deletions

Table S3 Structural Variants

Table S4 Discordant Uniquely Mapped Read Pairs

Table S5 Mass Spectrometry

Table S6 Mouse RNAseq

## Acknowledgements

We thank Daniel Bellott and other members of the Page and Mello laboratories for advice and discussion. We thank Eric Spooner at the Whitehead Proteomics Facility for mass spectrometry. We thank Kyomi Igarashi for technical assistance. We thank Peter Boag and Roger Pocock for access to microscopes and for technical advice. We thank Jurg Bahler for kindly allowing graphical representation of his yeast cell cycle data. Some strains were provided by the Caenorhabditis Genetics Center supported by NIH (P40 OD010440) and the International *C. elegans* Gene Knockout Consortium. This work was supported by Life Sciences Research Foundation to MAC; American Cancer Society 129916-PF16-232-RMC to GAD; NIH Grants (GM058800 and HD078253) to CCM. DCP and CCM are Howard Hughes Medical Institute Investigators.

## Author Contributions

Conceptualization, MAC, GAD; Investigation, MAC, GAD, GMD, TEG, LWM, HRT, DGD, KAR; Writing-Original Draft, MAC; Writing-Review and Editing, MAC, GAD, GMD, CCM; Funding acquisition, MAC, GAD, GMD, DCP, GJH, CCM; Supervision, MAC, DCP, GJH, CCM.

## Declaration of Interests

The authors declare no competing interests.

## METHODS

### CONTACT

Further information and requests for resources and reagents should be directed to and will be fulfilled by Michelle A. Carmell and Craig C. Mello.

### EXPERIMENTAL MODEL AND SUBJECT DETAILS

The N2 Bristol strain of *C. elegans* was cultured at 20°C under standard conditions as described in Brenner (Brenner, 1974). Deletion alleles of *gcna-1* and *dvc-1* were generated using RNP/*rol-6* strategy (Dokshin et al., 2018), outcrossed to N2, and balanced with nT1[qls51] or qC1[qls26].. The nature of the *gcna-1* alleles (on LGIII) is as follows: *ne4444*: 6006586/6006587-6008976/6008977 (deletion of entire coding sequence with a small insertion inside the breakpoints (AAATTCCTAAAATTTCCTGTATTC)); *ne4356*: 6007278/6007279– 6009026/6009027 (1748-bp deletion, removes ATG) (Carmell et al., 2016). The *dvc-1(ne4443)* deletion allele on LGV deletes the entire coding sequence: ChrV: 11237535/6-11238944/45 (deletion of entire coding sequence). The *dvc-1(ok260)* allele was obtained from the CGC (Strain ID RB1401). The fusion protein lines (*gfp::gcna-1, top-2::mcherry*, and *mcherry::dvc-1*) were made by CRISPR with hybrid donors as described in (Dokshin et al., 2018).

*Gcna*-mutant mice are deposited at the Jackson Laboratory, Bar Harbor, ME.

### METHODS DETAILS

#### *C. elegans* immunohistochemistry

Whole mount preparations of dissected gonads, fixation, and immunostaining procedures were carried out as described in (Dawson et al., 2017). Both PGL-1 (gift from Peter Boag) and PH3 (Merck Millipore) antibodies were used at 1:300 dilutions. Secondary antibodies and DAPI (Abcam) were applied at 1:1000. Images were observed using a Zeiss Axio Imager M2 and captured using an Axiocam 506 mono camera (Zeiss). Figures were constructed using Photoshop and Illustrator (Adobe) and graphs and statistical analysis was performed using Graphpad Prism (Graphpad).

#### *C. elegans* topoisomerase inhibitor, UV and drug treatments

Inhibition of topoisomerase, DNA replication, and inducing dsDNA breaks was achieved by subjecting worms to plates prepared with 70μM etoposide (Sigma-Aldrich), 25mM hydroxyurea (Sigma-Aldrich), and 50μM camptothecin (Sigma-Aldrich) respectively and were performed in triplicates three times as described previously (MacQueen and Villeneuve, 2001). In short, twenty L4 staged worms of wildtype and mutant strains were placed on NGM plates enriched with each poison, seeded with OP50 and incubated at 20°C and 25°C for 16 h. Worms were then transferred to seeded NGM plates with no poisons for 4 h for recovery, then removed. Plates with embryos were then incubated at their respective temperatures for 24 h after which time hatching rates were determined.

#### *C. elegans* acridine orange staining

Germ cell undergoing apoptosis were assessed *in vivo* via acridine orange as described previously (Boag et al., 2005). Briefly, 20 young adult worms were placed on NGM plates seeded with OP50 and stained with 1ml of 100μM acridine orange (Sigma-Aldrich) for 1 hour, then washed in M9 buffer and immobilized on 2% agarose gel pads in 0.03% tetramisole and observed using DIC and fluorescence microscopy. Each assay was performed at 20°C and 25°C in duplicates and repeated 3 times.

#### *C. elegans* mutator assay

*unc-58(e665)* mutator assay was carried out as in (Harris et al., 2006). Briefly, thirty 6cm plates were seeded with OP50 and 5 worms doubly mutant for either *gcna-1* or *dvc-1* and *unc-58(e665)* were added and incubated at 20°C. After several generations, the entirety of each plate of starved worms was chunked onto a large 10cm plate with concentrated OP50 on one side. Plates were scored one week later for revertant worms that were no longer small and paralyzed.

#### *C. elegans* genomic DNA sequencing

After isolating genomic DNA from worm strains, libraries were prepared using NEBNext^®^ Ultra™ DNA Library Prep Kit for Illumina and sequenced on an Illumina NextSeq machine using a TG NextSeq^®^ 500/550 Mid Output Kit v2 (150 cycles).

#### Bioinformatics

Sequence data were demultiplexed and adapter sequences were trimmed from reads using the FASTQ Generation workflow on the Illumina BaseSpace Sequence Hub. Sequence reads were further pre-processed using fastp v0.19.6 (Chen et al., 2018) to trim poly-G tails and aligned to the C. elegans reference genome assembly WBcel235 using BWA MEM v0.7.17 (Li, 2013). Sequence data from 4 lanes of each of 2 runs, one with 37 – 38 nucleotide paired end reads and the other with 150 nucleotide paired end reads, were merged into single BAM files for each sample. Duplicate read pairs based on aligned positions of each end were marked using Picard v2.18.12 (http://broadinstitute.github.io/picard). Alignment metrics were computed using Picard CollectAlignmentSummaryMetrics, CollectInsertSizeMetrics and CollectWgsMetrics. Poorly mapped regions for which over 10% of aligned reads are ambiguously placed, multi-mapping reads were determined using the CallableLoci tool from GATK v3.8.0 (McKenna et al., 2010).

Copy number analysis was carried out using VarScan v2.4.3 (Koboldt et al., 2012), CNVnator v0.3.3 (Abyzov et al., 2011) and Control-FREEC v11.5 (Boeva et al., 2012) using the unc-58 sample as a control (note there was a lower mapping rate and hence lower sequencing depth in the N2 control due to likely bacterial contamination). Circular binary segmentation was performed on the relative copy number computed by VarScan using DNAcopy v1.54 (Olshen et al., 2004)]. Homozygous deletions called by VarScan were assessed using the Integrative Genomics Viewer (IGV) (Robinson et al., 2011).

Genomic rearrangements in each of the 6 mutant samples that are not present in either of the parental strains (N2 and unc-58) were identified using three structural variant callers: Manta v1.5.0 (Chen et al., 2016), SvABA v0.2.1 (Wala et al., 2018) and Pixie v0.6, an in-house discordant read pair and split read clustering tool. Consensus structural variant calls made by and passing filters applied by at least 2 of the 3 callers were assessed using IGV.

#### *C. elegans* mortal germline assay

Mortal germline assays were performed at 25°C where 10 Individual L1 wild-type and *gcna-1* mutants were placed on individual seeded agar plates until they laid progeny. One worm from the progeny of each plate were transferred to new plates and allowed to mature. This process was repeated until worms were sterile. Brood size and rates of embryonic lethality assays were conducted on each generation.

#### *C. elegans* live cell imaging

Young adults were dissected on slides in M9 to release early embryos. Embryos were collected and immediately mounted on agar pads for imaging. Imaging was carried out using Zeiss Axio Imager M2 (Zeiss) with images collected every 15 to 30 seconds. Brightness and contrast were adjusted in Adobe Photoshop (Adobe). Brightness in some panels of Figure 3B was increased relative to earlier images in the time course to compensate for bleaching of the fluorescent signal over time, but does not affect the interpretation of this qualitative data.

#### *C. elegans* co-immunoprecipitation

Frozen pellets of 100,000 synchronized gravid adults were broken by bead beating in 1.5X lysis buffer containing 250 mM Tris HCl pH 7.5, 150mM Sodium Chloride, and 50mM Sodium Citrate (supplemented with protease inhibitors). The lysate was sonicated on ice at 30% amplitude for 3 minutes (15 seconds on 45 seconds off) followed by 40% amplitude for 30 seconds (15 seconds on 45 seconds off) in a Bioruptor (Diagenode). The sonicated lysate was supplemented with 1% NP-40 alternative and incubated rotating for 1 hour at 4C. Carcasses were spun down and supernatant was pre-cleared with 100uL pre-washed Protein G Dynabeads rotating at 4C for 1 hour. Immunoprecipitations were carried out at 4C overnight with mouse anti-GFP monoclonal antibody (Wako 018-20463), mouse anti-mCherry monoclonal [1C51] (Abcam ab125096) or control mouse IgG. Antibody was captured with 100ul pre-washed Protein G Dynabeads at 4C for 2 hours, washed 3x with wash buffer containing 250 mM Tris HCl pH 7.5, 150mM Sodium Chloride, 250mM Sodium Citrate, and 0.5% NP-40 alternative (supplemented with protease inhibitors). Protein was eluted of the beads for 10 minutes at 50°C in 50 ul of 1X NuPAGE™ LDS Sample Buffer (Invitrogen NP0007) with 100mM DTT and 30ul of the elution was used for a western blot. Western blotting was performed with rabbit anti-mCherry polyclonal antibody (Abcam ab183628).

#### *C. elegans* RNAi

*chk-1* RNAi was done by feeding with clone V-13G06 and empty vector from the Ahringer RNAi Library (Source Bioscience) (Timmons et al., 2001). The N2 strain was used as wildtype in all experiments. IPTG-containing RNAi plates were seeded with cultures of HT115(DE3) carrying the appropriate vector. For RNAi treatment, L4 larvae were placed on RNAi plates for 20 hours at 20°C, and single picked onto blank plates for egg laying. Immediately after removal of the adult worm, embryos were counted and scored for hatching 24 hours later.

#### *C. elegans* brood counts

Brood and male frequency counts were performed at 20 and 25°C. Briefly, animals were single picked at mid-L4 stage and followed with daily transfers until they produced no more progeny. Animals were counted when the population on a progeny plate reached adulthood.

#### Phylogenetic tree construction

Alignments were generated using MUSCLE (RRID:SCR_011812) (Edgar, 2004). ProtTest (RRID:SCR_014628) was used to select the best-fit model for protein evolution. PhyML (RRID:SCR_014629) version 20111216 was used to create maximum likelihood trees (Guindon et al., 2010). R6T986_9STAP and A0A0F7RMY7_BACAN, zinc metallopeptidases in the same Clan (CL0126), but a different family (PF01863), as GCNA were used as an outgroup.

#### Mouse testis histology

Mouse testes were fixed overnight in Bouin’s fixative at 4°C, then transferred to 70% ethanol before processing and embedding in paraffin. Five-micron sections were stained with hematoxylin and eosin before histological examination.

#### Mouse embryonic stem cell fluorescence activated cell sorting

Embryonic stem cells were immunostained with anti-GCNA antibody and a FITC labeled secondary antibody, stained with propidium iodide, and analyzed by flow cytometry using a FACS Aria II sorter (BD Biosciences) using BD FACSDiva Software Version 8.0 and FlowJo Software Version 10.5.3. Levels of GCNA expression were binned such that 51.2% of cells fell in the “GCNA medium” category.

#### Calculation of crossover interference

To estimate the strength of crossover interference, we fit a gamma distribution to the distances between MLH1 foci, following the method of (de Boer et al., 2006). Briefly, we binned inter-crossover distances by chromosome length, and fit the observed distribution of inter-crossover distances to a gamma distribution using Scipy, obtaining initial estimates of the gamma shape and scale parameters. Then, we refined our shape and scale estimates, used a simulation approach to correct for the fact that interfocus distances cannot be greater than chromosome length or shorter than the resolution of our immunofluorescence images.

#### RNA-seq (gene expression and transposon analysis)

Total RNA was isolated from the testes of 2 wildtype and 2 *Gcna*-mutant mice using Trizol at both postnatal day 8 and postnatal day 18. RNA was enriched for polyA and sequencing libraries were prepared by the Whitehead Institute Genome Core Facility and sequenced on an Illumina HiSeq with 75 bp paired end reads. For genome-wide differential expression analysis, reads were aligned to the mouse genome (mm10) using TopHat and fold-changes and p-values were calculated using cuffdiff (Trapnell et al., 2012). For analysis of transposon expression, LTR and non-LTR retrotransposon sequences were downloaded from repBase (Bao et al., 2015), reads were aligned to the retrotransposon sequences with bowtie2 (Langmead et al., 2009), expression of each transposon was quantified with eXpress (Roberts and Pachter, 2013), and differential expression analysis was performed with edgeR (Robinson et al., 2010).

#### Mouse spermatocyte spreads

Mouse spermatocyte spreads were carried out as in (Peters et al., 1997). Meiotic cells were isolated from mascerated seminiferous tubules, spun down and resuspended in hypotonic buffer (30 mM Tris-HCl pH 8.2, 50 mM sucrose pH 8.2, 17 mM sodium citrate). After a second spin they were resuspended in 0.1 M sucrose and dropped onto the slides wet with 1% PFA, 0.1% Triton X-100 in sodium borate buffer pH 9.2 and incubated in a humid chamber for 2–3 hr. For immunostaining, slides were blocked in 3% BSA and incubated with primary and secondary antibodies in 1% BSA. Nuclei were stained using the following antibodies: mouse anti-H2A.X phosphorylated on Ser 139 (anti-γH2A.X) (Abcam),rabbit anti-γH2A.X (Abcam), goat anti-ATR (Santa Cruz), rabbit anti-mouse BRCA1 (gift of S. Namekawa), mouse anti-SYCP3 (Santa Cruz), mouse anti-MLH1 (Millipore). Images were collected using a DeltaVision system (Applied Precision) and subjected to deconvolution and projection using the SoftWoRx 3.3.6 software (Applied Precision).

#### Embryonic stem cell survival assay

ES cells were cultured under standard conditions. Cells were trypsinized, counted, and 500 cells were plated at clonal density in triplicate for each condition. After cells had adhered to the plate, media was removed and cells were irradiated with indicated doses of UV using a Stratalinker. 7-10 days later, surviving colonies were stained with Crystal violet and counted.

#### Immunoprecipitation from mouse embryonic stem cells and mass spectrometry

Mouse embryonic stem cells were irradiated with 8J/m2 UV, harvested 1 hour later by scraping, and suspended in IP buffer: 50 mM HEPES pH 7.4, 140 mM NaCl, 10% glycerol, 0.5% NP-40, 0.25% Triton, 100 uM ZnCl2 plus EDTA-free Protease Inhibitor Tabs (Roche). Samples were sonicated on ice 1 min at 30% amplitude using a Branson Sonifier and treated with 100 U/ml Benzonase (Millipore) for 20 min at room temperature on rocker. Samples were spun down for 10 min at 16,000 G at 4°C, and supernatant was used for immunoprecipitations. After extensive washing in IP buffer, precipitated proteins were subjected to SDS–PAGE and silver staining. Samples were processed at the Whitehead Institute Proteomics Core Facility. For mass spectrometry analysis, bands were excised from each lane of a gel encompassing the entire molecular weight range. Trypsin digested samples were analyzed by reversed phase HPLC and a ThermoFisher LTQ linear ion trap mass spectrometer. Peptides were identified from the MS data using SEQUEST (RRID:<u>SCR_014594</u>).

## Supplemental Information

Table S1: alignment_metrics

Alignment metrics for merged whole genome sequencing data including the total numbers of read pairs, the percentage of those that align to the WBcel235 reference genome, the median insert size (inferred fragment length), the percentage of read pairs that are marked as duplicates based on aligned positions of both ends, the median coverage and the percentage of bases in the reference genome that reach various depths of coverage. All metrics were computed using the Picard tools: CollectAlignmentSummaryMetrics, MarkDuplicates CollectInsertSizeMetrics and CollectWgsMetrics.

Table S2: homozygous_deletions

Homozygous deletions called by VarScan.

Table S3: structural_variants

Consensus structural variant calls made by at least 2 of Manta, SvABA and Pixie. The breakpoints for each junction end are given as a range with the Chromosome, Start and End columns. Variants supported by breakpoint-spanning reads with split read alignments are typically resolved to single base pair resolution.

Manta filters:

MinSomaticScore: Somatic score < 30

SvABA filters:

COMPETEDISC: Discordant cluster found with nearly the same breakpoints but differing strands

LOWAS: Alignment score of one end is less than 80% of the contig length or the number of mismatch bases on one end >= 10

LOWMAPQDISC: Both clusters of reads failed to achieve a mean mapping quality > 30 NODISC: Rearrangement was not detected independently of assembly

WEAKDISC: Fewer than 7 supporting discordant reads and no assembly support

Pixie filters:

SupportInControl: Fraction of total supporting read pairs from control samples > 0.05

Table S4: discordant_uniquely_mapped_read_pairs Table S5: mass_spectrometry

Table S6: mouse_rnaseq

Figure S1: Gcna-1 mutant germ cells exhibit normal proliferation, RNA granule morphology, and slightly elevated apoptosis relative to wildtype.

Figure S2: Embryo hatching in the absence of drug treatment was not significantly different between strains.

Figure S3: Rearrangements at the *unc-58* locus in *gcna-1(ne4356);unc-58(e665)* revertants

Figure S4: *Gcna*-mutant ES cell survival is not affected by UV irradiation.

Figure S5: *Gcna*-mutant spermatocytes exhibit asynapsis, DNA damage, and chromatin condensation abnormalities.

Figure S6: Cumulative distribution curves of the distance between two MLH1 foci in wildtype and mutant bivalents (Chromosomes 11-13).

