## Supplementary Figures 1-6 for "GCNA interacts with Spartan and Topoisomerase II to regulate genome stability"

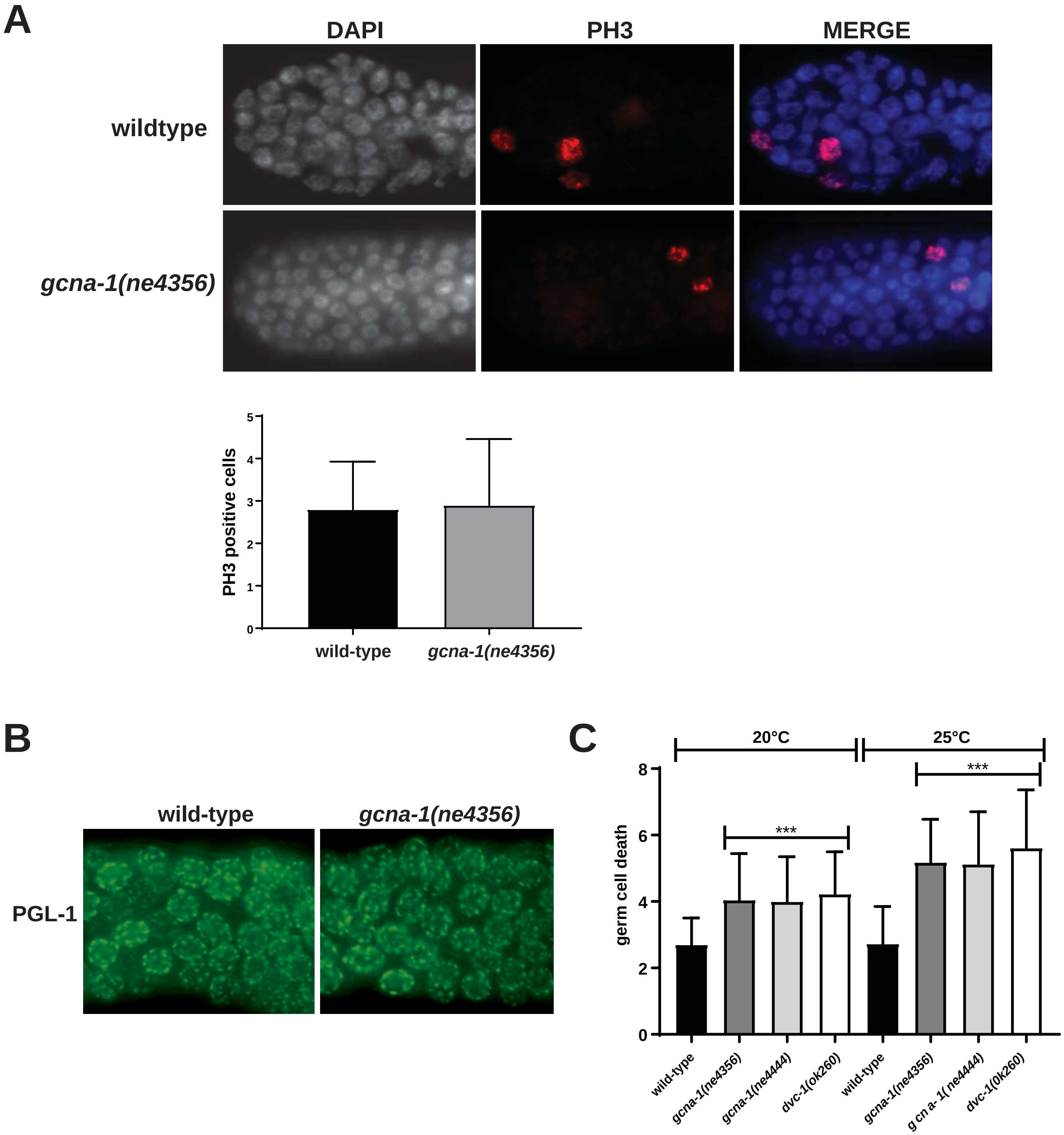


Figure S1: *Gcna-1* mutant germ cells exhibit normal proliferation, RNA granule morpholo-gy, and slightly elevated apoptosis relative to wildtype. A. Mitotic zones of wildtype and *gcna-1* mutant (ne4356) germlines immunostained with antibody recognizing phospho-histone H3 (PH3). B. Meiotic zone of gonads immunostained P-granule marker PGL-1. C. Quantification of germ cell death as assayed by acridine orange staining.

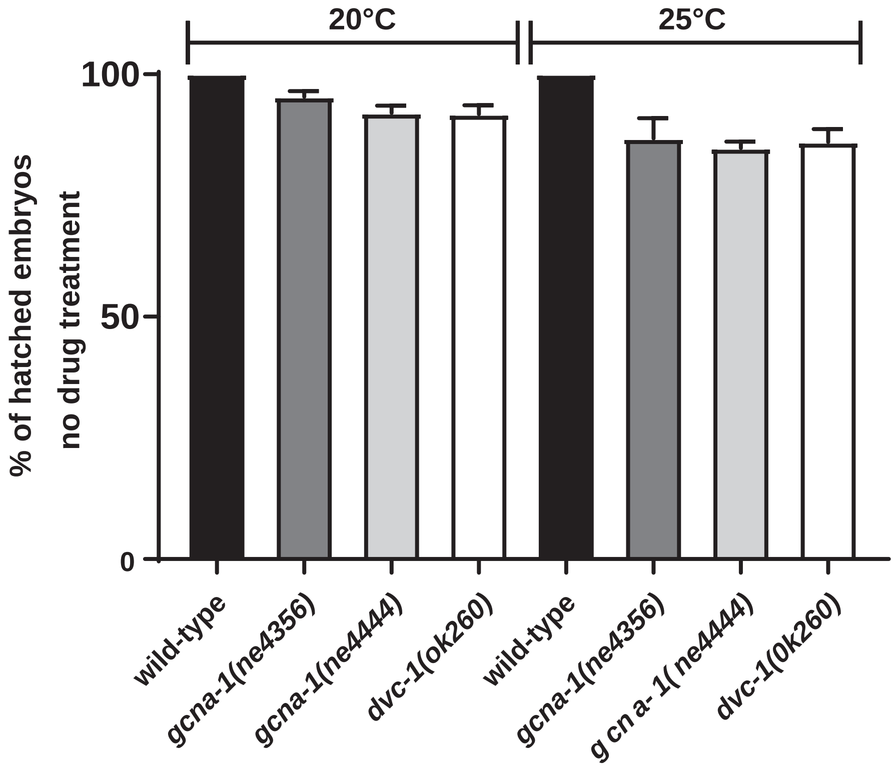


Figure S2: Embryo hatching in the absence of drug treatment was not significantly different between strains. Animals were treated as in drug exposure trials, but with no drug and hatching rates were assayed.

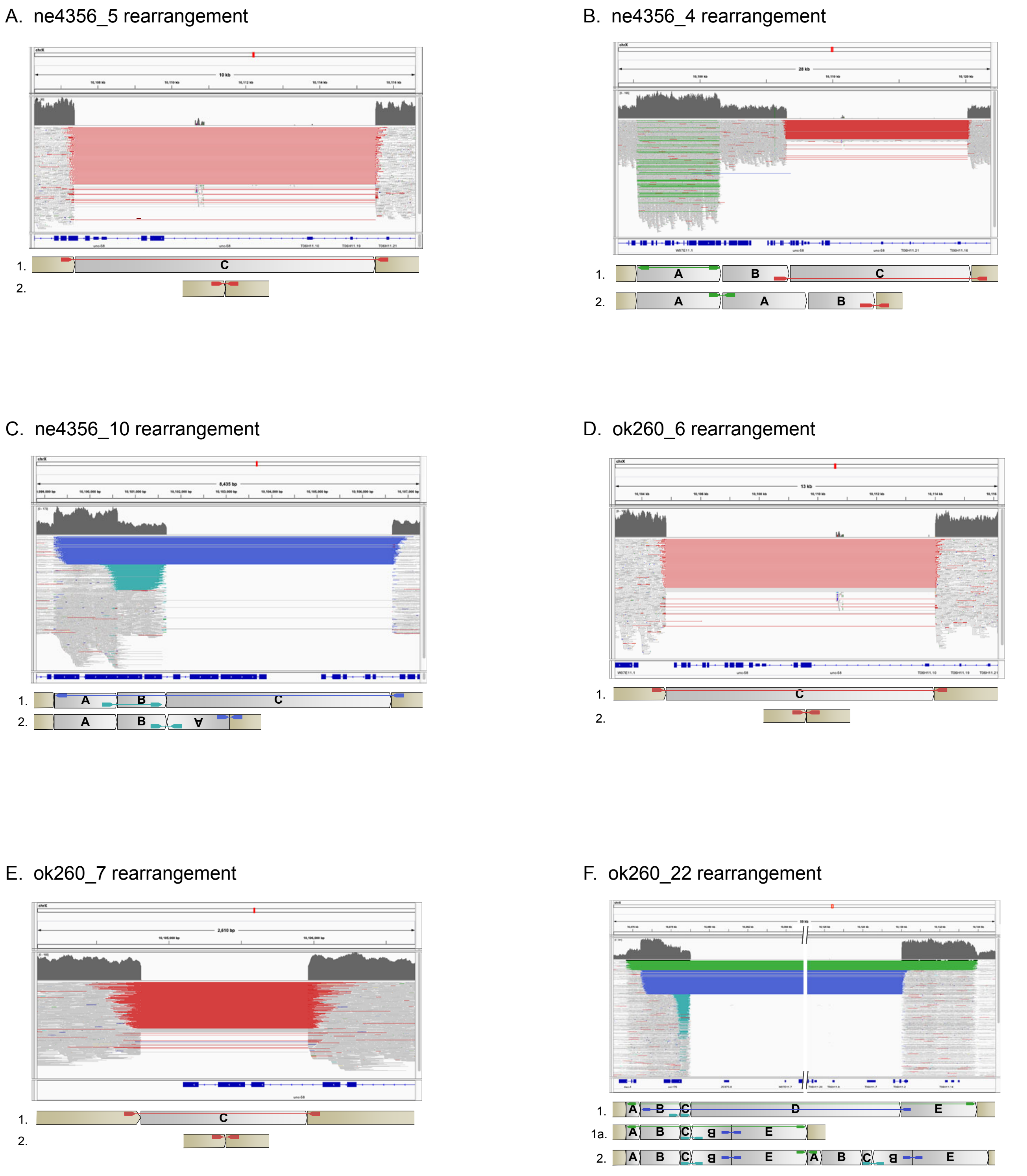


Figure S3: Rearrangements at the *unc-58* locus in *gcna-1(ne4356);unc-58(e665)* revertants (A-C) and *dvc-1(ok260);unc-58(e665)* revertants (D-F). For each rearrangement, Line 1 shows how discordant reads map on the reference sequence. Line 2 shows the rearrangement that can resolve the discordant reads. In panel F, Line 1a shows an intermediate rearrangement step that preceded a duplication of the rearrangement. Color coding of sequencing reads is according to the default settings in the Integrative Genomics Viewer (Broad Institute) and is as follows: Black, normal reads; Red, deletion (inferred insert size is larger than expected); Cyan, +/+ reads (implies inversion); Blue, -/- reads (implies inversion); Green, outward facing reads (implies duplication or translocation).

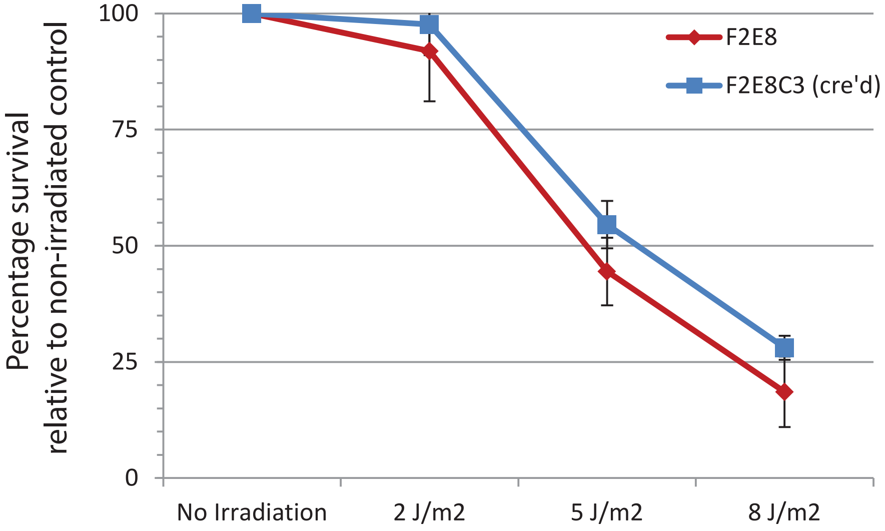


Figure S4: *Gcna*-mutant ES cell survival is not affected by UV irradiation. F2E8 is an ES cell line harboring Cre-recombinase sites flanking Exon 4 of the *Gcna* gene. F2E8C3 is the same cell line after treatment with Cre-recombinase, which deletes Exon 4, causing a frameshift affecting the remaining exons, drastically reducing the amount of mRNA transcribed from the *Gcna* gene. T-test; $P>0.1$ at all doses.

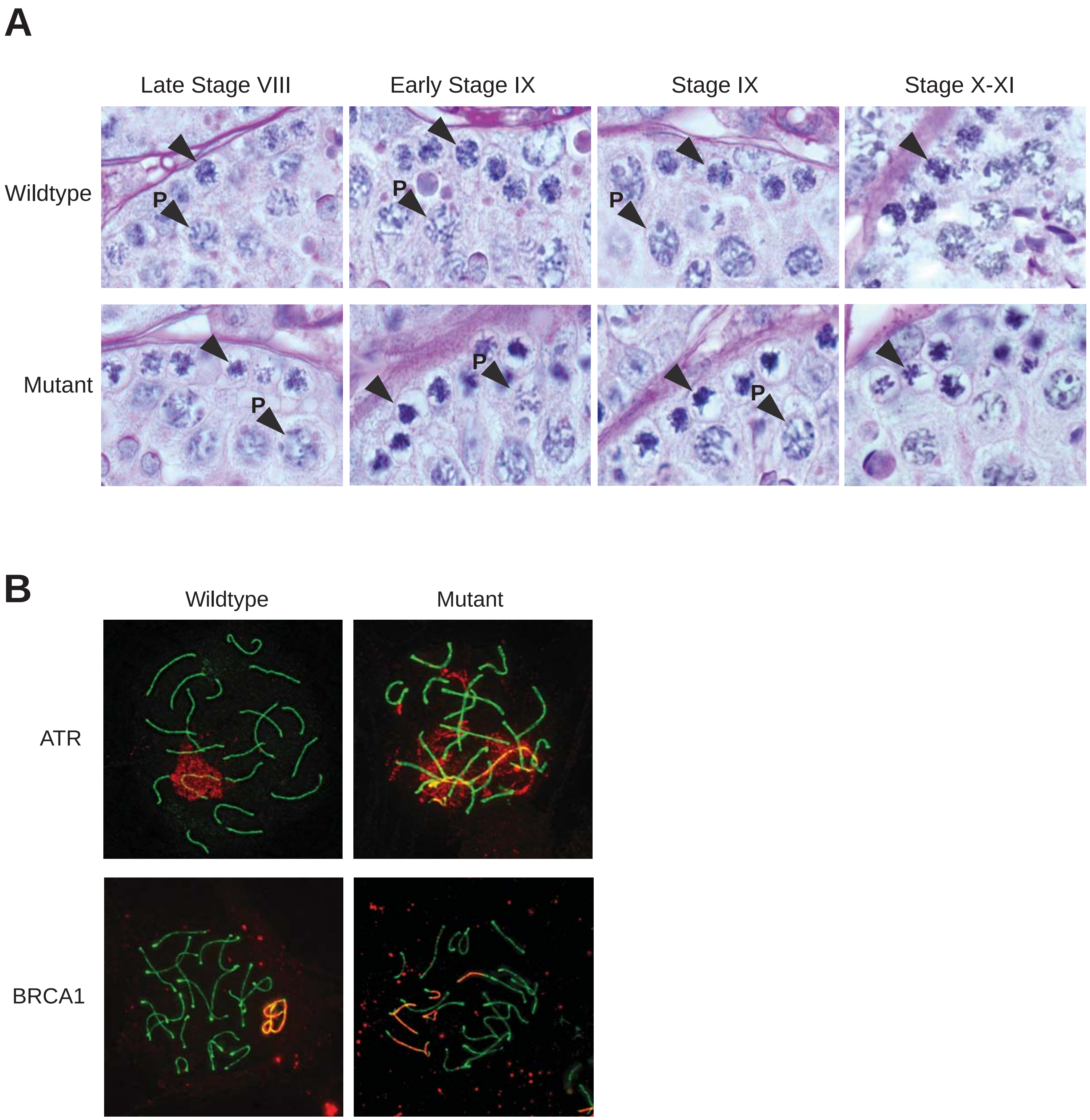


Figure S5: Gcna-mutant spermatocytes exhibit asynapsis, DNA damage, and chromatin condensation abnormalities. A. Aberrant chromatin condensation begins in the leptotene stage of prophase of Meiosis I. Wildtype and *Gcna* mutant hematoxylin and eosin stained mouse testis sections showing Stage VIII-XI seminiferous tubules. Representative leptotene (Stage VII-IX) and zygotene (X-XI) spermatocytes are indicated by arrows. Pachytene spermatocytes are indicated by "P". Note that the Stage IX sections are the same as those depicted in the main figure. B. Wildtype (*Gcna*$^{+/Y}$) and *Gcna* mutant (*Gcna*$^{DeltaEx4/Y}$) spermatocytes immunostained with antibodies to synaptonemal complex component SCP3 (green) and either unsynapsed chromosome marker BRCA1 (red) or DNA damage marker ATR (red) as indicated.

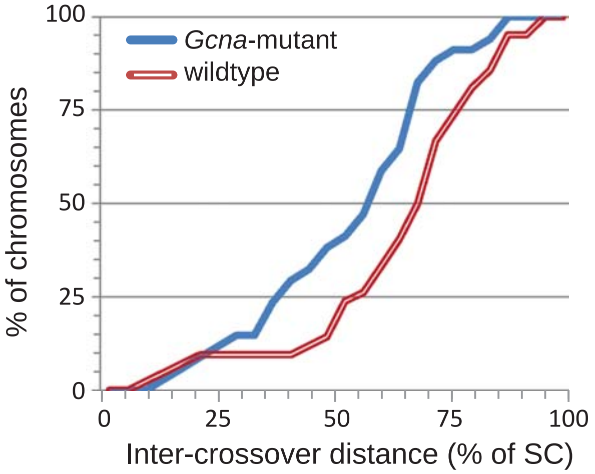


Figure S6: Cumulative distribution curves of the distance between two MLH1 foci in wildtype and mutant bivalents (Chromosomes 11-13). Crossover interference gamma shape parameter for wildtype is 11.037 and mutant is 9.479 (Two-sided Mann-Whitney; p=0.0074).
